# Single-cell Lineage Tracing by Integrating CRISPR-Cas9 Mutations with Transcriptomic Data

**DOI:** 10.1101/630814

**Authors:** Hamim Zafar, Chieh Lin, Ziv Bar-Joseph

## Abstract

Recent studies combine two novel technologies, single-cell RNA-sequencing and CRISPR-Cas9 barcode editing for elucidating developmental lineages at the whole organism level. While these studies provided several insights, they face several computational challenges. First, lineages are reconstructed based on noisy and often saturated random mutation data. Additionally, due to the randomness of the mutations, lineages from multiple experiments cannot be combined to reconstruct a consensus lineage tree. To address these issues we developed a novel method, LinTIMaT, which reconstructs cell lineages using a maximum-likelihood framework by integrating mutation and expression data. Our analysis shows that expression data helps resolve the ambiguities arising in when lineages are inferred based on mutations alone, while also enabling the integration of different individual lineages for the reconstruction of a consensus lineage tree. LinTIMaT lineages have better cell type coherence, improve the functional significance of gene sets and provide new insights on progenitors and differentiation pathways.

## Introduction

Reconstructing cell lineages that lead to the formation of tissues, organs and complete organisms is of crucial importance in developmental biology. Elucidating the lineage relationships among the diverse cell types can provide key insights into the fundamental processes underlying normal tissue development as well as valuable information on what goes wrong in developmental diseases (1; 2; 3). Traditionally, heritable markers have been utilized for prospective lineage tracing by first introducing them in a cell and then using them to track its descendants (3). Such studies resorted to using diverse markers such as viral DNA barcodes (4), fluorescent proteins (5), mobile transposable elements (6), *Cre*-mediated tissue-specific recombination (7) and more. Other methods relied on retrospective lineage tracing by using naturally occurring somatic mutations (8; 9), microsatellite repeats (10) or epigenetic markers (11). While these approaches provided valuable insights, they are often limited to a small number of markers and cells and due to the lack of coupled gene expression information, they cannot characterize the diverse cellular identities of the tracked cells and their relation to the lineage branching (1).

Recent advances in single-cell transcriptomics (scRNA-seq) allow the profiling of thousands of individual cells and the identification of cell types at an unprecedented resolution (12; 13; 14). Cost-efficient and scalable technologies provide large-scale scRNA-seq datasets that can be used to identify gene expression signatures of diverse cell types and to curate catalogs of cellular identities across tissues (13; 15; 16). While some of these datasets have been used to infer developmental lineages (17), methods for such inference rely heavily on strong assumptions regarding expression coherence between developmental stages, which may not hold in all cases (17; 18). Moreover, these approaches alone are unable to recover intermediate cell types and states making it difficult to reconstruct the early developmental lineages in an adult organism (18; 19).

Very recently, new experimental techniques that simultaneously recover transcriptomic profiles and genetic lineage markers from the same cell have been introduced (20; 21; 22). One of the earliest methods using such approach is scGESTALT (20) which combines the CRISPR-Cas9-based lineage tracing method termed GESTALT (23) with droplet-based single-cell transcriptomic profiling. scGESTALT inserts Cas9-induced stochastic (random) mutations to a genomic CRISPR barcode array at multiple time points. The edited barcodes are then sequenced (using scRNA-seq) and utilized for reconstructing a lineage tree based on maximum parsimony (MP) criterion (24). Cell types are independently inferred based on the scRNA-seq data. While this and similar methods have been successfully applied to a number of organisms (20; 21), they suffer from several problems. First, the random mutation data used for reconstructing the MP lineage is noisy and often saturated making it difficult to separate different cell types, especially at later stages. Even though expression information is collected for all genes in each cell, to date the reconstruction of the lineage tree solely depends on the stochastic Cas9-induced mutations. As a result, the resulting lineage tree sometimes fails to separate different types of cells and places similar cell types on distant branches. Further, multiple tree topologies can have the same parsimony score based on mutations making the reconstruction more challenging. In addition, the random nature of the induced mutations restricts the lineage reconstruction to each individual and mutation data from multiple individuals cannot be combined for inferring a consensus lineage tree based on multiple experiments.

To improve the reconstruction of lineages from CRISPR-Cas9 mutations and scRNA-seq data, we developed a novel statistical method, LinTIMaT (<u>**Lin**</u>eage <u>**T**</u>racing by <u>**I**</u>ntegrating <u>**M**</u>utation <u>**a**</u>nd <u>**T**</u>ranscriptomic data) that integrates mutational and transcriptomic data for reconstructing lineage trees in a maximum-likelihood framework. LinTiMaT employs a novel likelihood function for evaluating different tree structures based on mutation information. It then defines a new likelihood optimization problem which combines the likelihood score for the mutation data with Bayesian hierarchical clustering (25), which evaluates the coherence of the expression information such that the resulting tree concurrently maximizes agreement for both transcriptomics and genetic markers from the same cell. The tree space is explored by a novel heuristic search algorithm that first infers a lineage tree based on mutation information and further refines it based on both mutation and expression information. Finally, LinTiMaT also introduces an algorithm for integrating lineages reconstructed for different individuals of the same species for inferring a consensus lineage tree. We applied LinTIMaT to two zebrafish datasets generated using scGESTALT and show that by integrating transcriptomic and mutational data, the method was able to reconstruct lineages that properly explained the mutation data as well as preserved the cell type coherence in different subtrees. In addition, for the first time, data from multiple individuals studied using scGESTALT could be combined by LinTIMaT for reconstructing the consensus lineage that further improved on each of the individual lineages both in terms of clade homogeneity and in terms of functional assignment for the cells residing on leaves of the lineage tree.

## Results

### Overview of LinTIMaT

To enable the accurate reconstruction of individual and consensus lineages we developed LinTIMaT that integrates CRISPR-Cas9 mutations with transcriptomic data from single cells. An overview of the algorithm is shown in Fig. 1. We assume that the cell lineage tree is a rooted directed tree (Fig. 1a). The root of this lineage tree denotes the initial cells that do not contain any marker (or editing event). The leaves of this tree denote the cells from which the mutated barcodes and RNA-seq data have been recovered. LinTIMaT reconstructs the lineage tree by maximizing a likelihood function that accounts for both mutations and expression data. The likelihood function imposes a Camin-Sokal parsimony criterion for each synthetic marker. The probability associated with a transition of mutation state for a marker along a branch of the lineage tree is computed based on the abundance of the marker in the single cells. To compute the expression likelihood based on the transcriptomic data, the lineage is modeled as a Bayesian hierarchical clustering (BHC) (25) of the cells and the marginal likelihoods of all the partitions consistent with the given lineage tree are computed based on a Dirichlet process mixture model. To optimize the tree topology, we employ a heuristic search algorithm, which stochastically explores the space of lineage trees.

**Figure 1:**
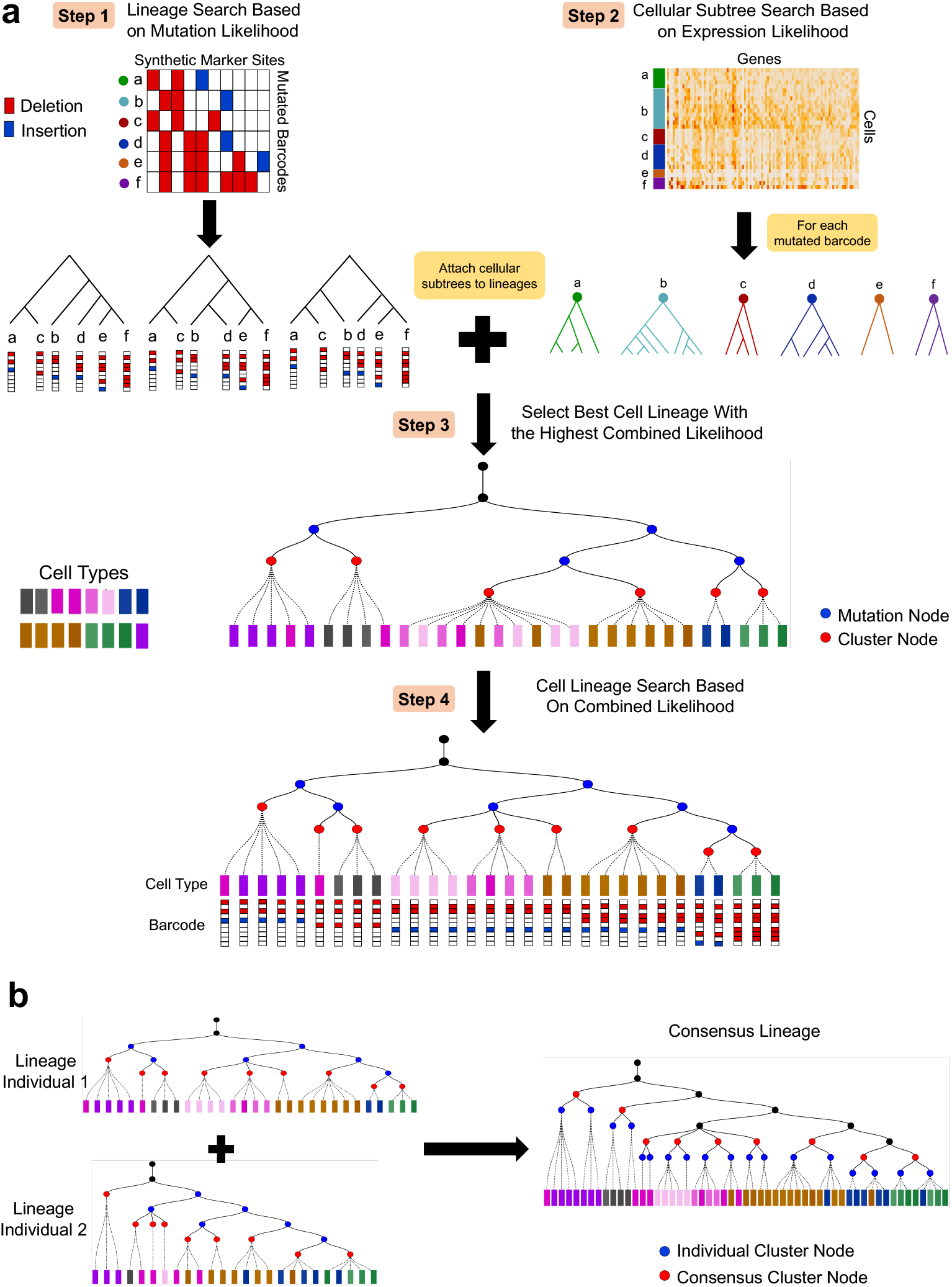
Overview of LinTIMaT. (a) LinTIMaT reconstructs a cell lineage tree by integrating CRISPR-Cas9 mutations and transcriptomic data. In Step 1, LinTIMaT infers top scoring lineage trees built on barcodes using only mutation likelihood. In Step 2, for all cells carrying the same barcode, LinTIMaT reconstructs a cellular subtree based on expression likelihood. In Step 3, cellular subtrees are attached to barcode lineages to obtain cell lineage trees and the tree with the best combined likelihood is selected. Finally, LinTIMaT uses a hill-climbing search for refining the cell lineage tree by optimizing the combined likelihood (Step 4). (b) To reconstruct a consensus lineage, LinTIMaT performs an iterative search that attempts to minimize the distance between individual lineage trees and the consensus tree topology. As part of the iterative process, LinTIMaT matches clusters in one individual tree to clusters in other individual tree(s) such that leaves in the

The above algorithm reconstructs trees for a specific CRISPR-Cas9 mutation set. To integrate trees resulting from repeat experiments of the same organism, LinTIMaT further reconstructs a consensus lineage tree (Fig. 1b). Our consensus lineage tree reconstruction method first infers cell clusters from the reconstructed individual lineages and then performs a greedy matching to pair the clusters from different individual lineages based on the similarity of gene expression data. Using this initial matching, it iterates to minimize an objective function consisting of two distance functions, the first is aimed at minimizing the disagreement between the topology of the consensus lineage tree and the individual lineage trees while the second distance is minimized for improving the cluster matching. See Methods for complete details.

### Integration of mutation and transcriptomic data improves the reconstruction of cell lineage trees

We applied LinTIMaT on two zebrafish datasets (ZF1 and ZF3) generated using scGESTALT (20). ZF1 and ZF3 consisted of 750 and 376 cells respectively, from which both the transcriptome (20287 genes) and edited barcode (192 unique barcodes, 324 unique markers for ZF1 and 150 unique barcodes, 265 unique markers for ZF3) were recovered. For both datasets, our analysis shows that improving the likelihood function used by LinTIMaT increases the coherence of the resulting cell types for each subtree, without impacting the overall mutation likelihood (Fig. 2a and Supplementary Fig. S1). For both fishes, LinTIMaT generated highly branched multiclade lineage trees (Fig. 2b and Supplementary Fig. S2). Blue nodes on the tree represent mutation events assigned while red nodes represent the clusters identified based on gene-expression data. It is important to note that cluster nodes do not necessarily represent common ancestors for the cells underneath, instead, cluster nodes are just a way of grouping nearby cells together based on expression information without affecting the mutational ancestor-descendant relationships. ZF1 lineage tree comprised 25 major clades (level 1 tree nodes) and 113 cluster nodes, 77 of which consisted of more than one cell. ZF3 lineage tree comprised 17 major clades and 42 cluster nodes, 33 of which consisted of more than one cell. We compared the lineage trees reconstructed by LinTIMaT to the trees reconstructed using maximum parsimony (MP) as used in the original study (20) by comparing the accuracy of cell clusters in the trees. In the original study, 63 transcriptionally distinct cell types were identified using an unsupervised, modularity-based clustering approach from 6 zebrafish samples. We used this clustering to compute the Adjusted Rand Index (ARI) for the cell clustering obtained from a lineage tree (Methods). For MP lineage trees, the unique barcodes represent cell clusters as mutation information was the only basis for reconstructing the tree. For each fish, the lineage tree reconstructed by LinTIMaT resulted in better cell clustering (37.5% and 36.4% improvement in ARI for ZF1 and ZF3 respectively) compared to MP results based on mutation data alone (see Supplementary Table S1 and Supplementary Results for details).

**Figure 2:**
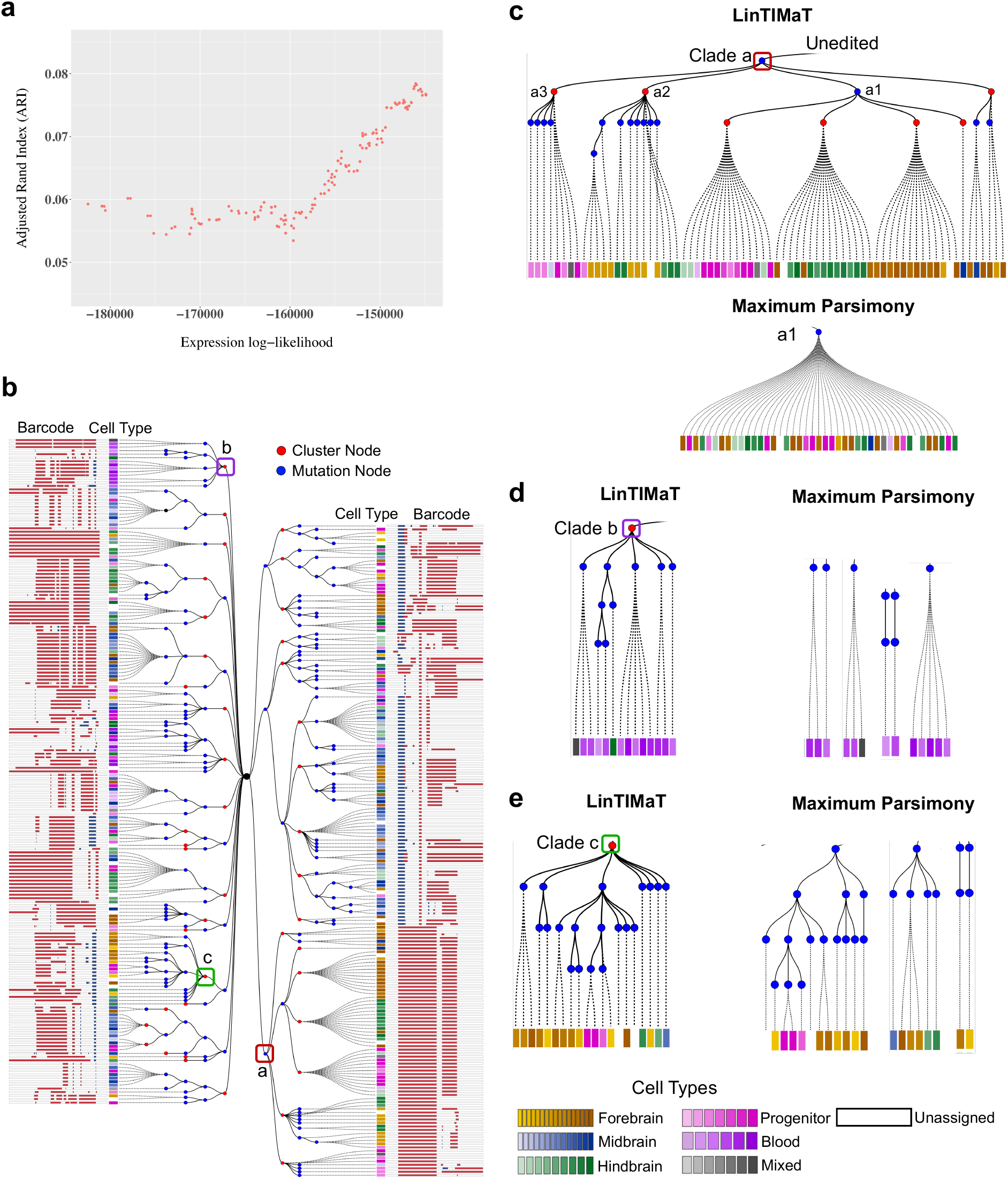
Reconstructed cell lineage for a single juvenile zebrafish brain (ZF3). (a) Adjusted Rand Index (ARI) as a function of the expression likelihood score calculated by LinTIMaT. The fact that both improve suggests a good agreement between the resulting tree and cell types. (b) Reconstructed cell lineage tree for ZF3 built on 376 cells. Blue nodes represent Cas9-editing events (mutations) and red nodes represent clusters inferred from transcriptomic data. Each leaf node is a cell, represented by a square, and its color represents its assigned cell type as indicated in the legend. The mutated barcode for each cell is displayed as a white bar with insertions (blue) and deletions (red). (c) By using transcriptomics data LinTIMaT is able to further refine subtrees in which all cells share the same barcode which can help overcome saturation issues. (d-e) Example subtrees displaying LinTIMaT’s ability to cluster cells with different barcodes together based on their cell types. In contrast, maximum parsimony puts these on distinct branches.

Lineage trees reconstructed using LinTIMaT showed successful integration of mutation and expression data. When using only mutation data, in several cases, cells belonging to very different cell types were clustered together. In contrast, in the trees reconstructed by LinTIMaT, these cells were correctly assigned to different subtrees corresponding to different cell types. Clade a1 in ZF3 lineage tree (Fig. 2c) is one such example. In MP lineage tree for ZF3, neural progenitor cells, hindbrain granule cells, and neurons in ventral forebrain and hypothalamus (total 43 cells) were clustered together under clade a1 as they shared the same mutational barcode. The tree reconstructed by LinTIMaT corretly separated these cells into three major subtrees (progenitor, hindbrain, and forebrain) under the same mutational node. Similarly for ZF1, in the original MP lineage tree, clade a consisted of 198 cells including mostly forebrain and progenitor cells. LinTIMaT lineage tree successfully divided them into multiple subtrees, with the largest mainly containing forebrain neuron cells and the other subtrees mostly containing different types of progenitor cells (Supplementary Fig. S3a). In addition, LinTIMaT trees also contain examples where cells belonging to similar cell types but carrying different mutational barcodes are identified as a cluster instead of being placed on distant branches as done by MP. Clades b and c in ZF3 lineage tree (Fig. 2d-e) illustrate this scenario. In the LinTIMaT lineage tree for ZF3, clade b consists of mostly blood cells that carry different mutational barcodes. In MP lineage tree, these cells were placed in 4 distant branches which did not convey the fact that they belong to the same cell type. However, LinTIMaT successfully grouped them together in a cluster of blood cells while preserving their mutational differences as illustrated by the mutation nodes being descendants of the cluster node. Similarly, for clade c most of the cells were forebrain neurons that were placed in three distinct branches in the MP lineage tree owing to their mutational differences. LinTIMaT successfully identified these cells as a cluster consisting of mostly forebrain neuron cells. Similar examples can be seen in the tree reconstructed by LinTIMaT for ZF1 (Supplementary Fig. S3b). We note that while the LinTIMaT reconstructed lineage trees displayed much better agreement with cell type coherence, this was not just a function of ignoring mutational data. In fact, the trees inferred by LinTIMaT have *higher* likelihoods based on mutation alone (Supplementary Table S2) when compared to the trees reconstructed by MP (20). In fact, for each fish, the MP lineage tree violated the Camin-Sokal parsimony criterion for some mutations that resulted in a low mutation log-likelihood.

Following the analysis of (20), we also analyzed the trees for spatial enrichment of clusters. For this, groups of four or more cells were selected for both LinTIMaT and MP lineage trees. In both types of lineage trees, clusters were spatially enriched in hindbrain, forebrain and midbrain (Fig. 3). However, the trees reconstructed by LinTIMaT displayed better spatial enrichment. For example, for ZF3, more clusters in LinTIMaT lineage tree were enriched in forebrain and hindbrain compared to the barcode clusters in MP tree. Similarly, for ZF1, LinTIMaT lineage showed more enriched hindbrain clusters compared to the barcode clusters in MP tree.

**Figure 3:**
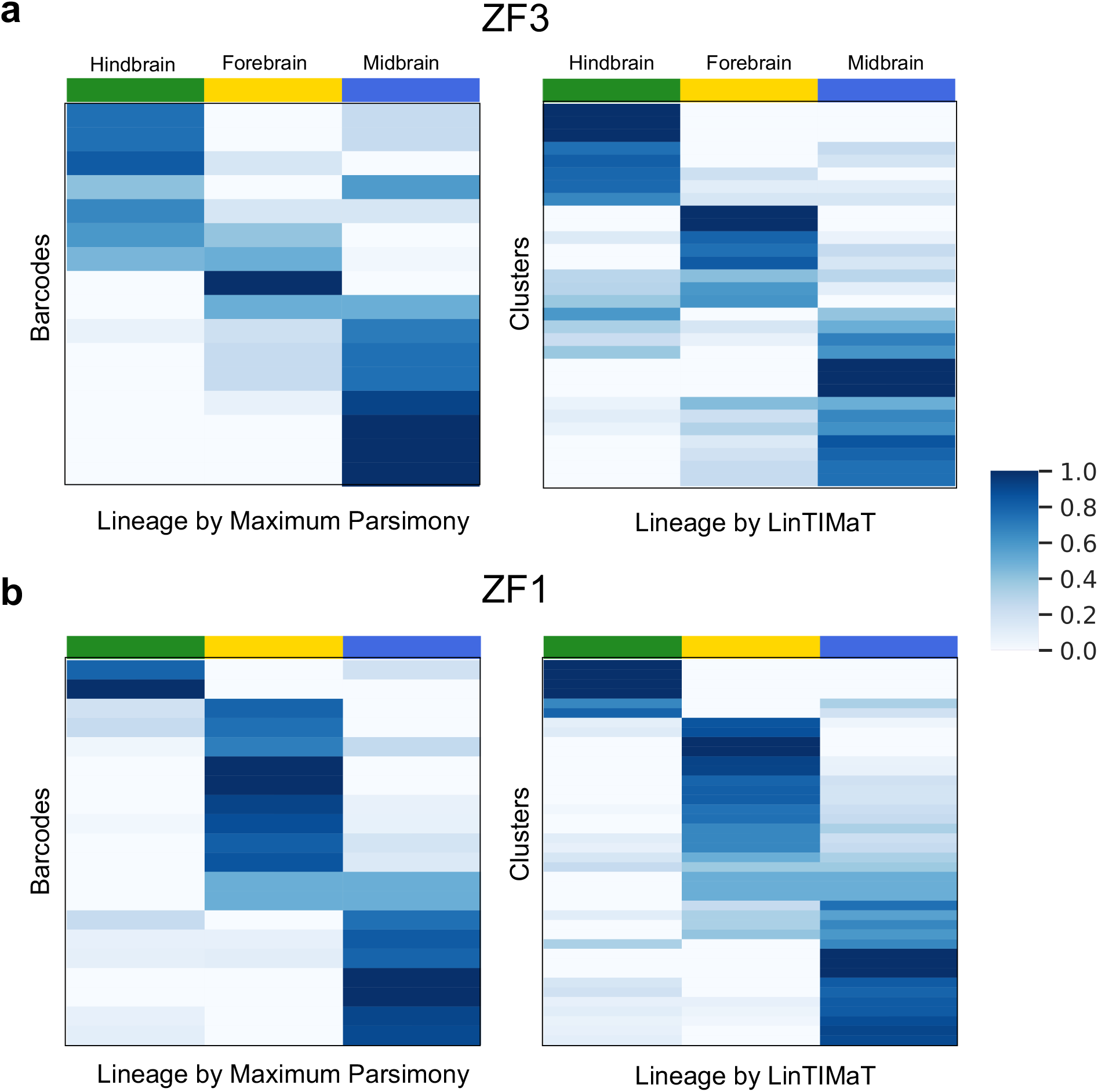
Distribution of cell types in the juvenile zebrafish brain. Heat map of the distribution of cell clusters for each region of the brain (columns). Cell types were classified as belonging to the forebrain, midbrain or hindbrain, and the proportions of cells within each region were calculated for each cluster. For MP lineage, the rows of the heat map represent barcodes, for LinTIMaT lineage, the rows represent clusters inferred from barcodes and expression data. (a) Comparison for ZF3. (b) Comparison for ZF1.

LinTIMaT lineage trees also revealed divergent lineage trajectories. For example, for ZF3, LinTIMaT lineage tree displayed three major subtrees under clade a (a1, a2 and a3 respectively), with a1 being sub-divided into three major clusters. Clade a1 had three major clusters consisting mostly of progenitor cells, hindbrain and forebrain neurons respectively. The constructed tree indicates that the *her*4.1^+^ and *atoh*1*c*^+^ progenitor cells (26; 27) are closely related to *pax*6*b*^+^ granule cells (28) in hindbrain, *gad*2^+^ neurons in ventral forebrain (29), and *fezf* 1^+^ neurons (30) in hypothalamus region. On the other hand, *pitx*2^+^ and *prdx*1^+^ neurons (31) in forebrain (clade a2) were determined to be related to radial glia cells (clade a3). These results demonstrate LinTIMaT’s ability to elucidate complex lineage relationships of cells.

### Consensus lineage tree successfully combines data from individual lineages

As mentioned above, combining CRISPR-Cas9-mutation-based individual lineage trees is challenging since mutations are random and so differ for the same cell types between experiments. To address this, we used LinTIMaT to combine data from both ZF1 and ZF3 in order to infer a consensus lineage for the development of juvenile zebrafish brain. Since the two fishes had a different number of cells, we subsampled 380 cells for ZF1 so that both fish have equal weights when learning the consensus tree. LinTIMaT inferred 43 clusters for ZF1 and 42 clusters for ZF3 and so we split one cluster in ZF3 to obtain 43 clusters for both trees (Methods). Using these, LinTIMaT inferred a consensus lineage tree (Fig. 4) with 43 leaves each of which represents a matched pair of clusters from the individual fishes.

**Figure 4:**
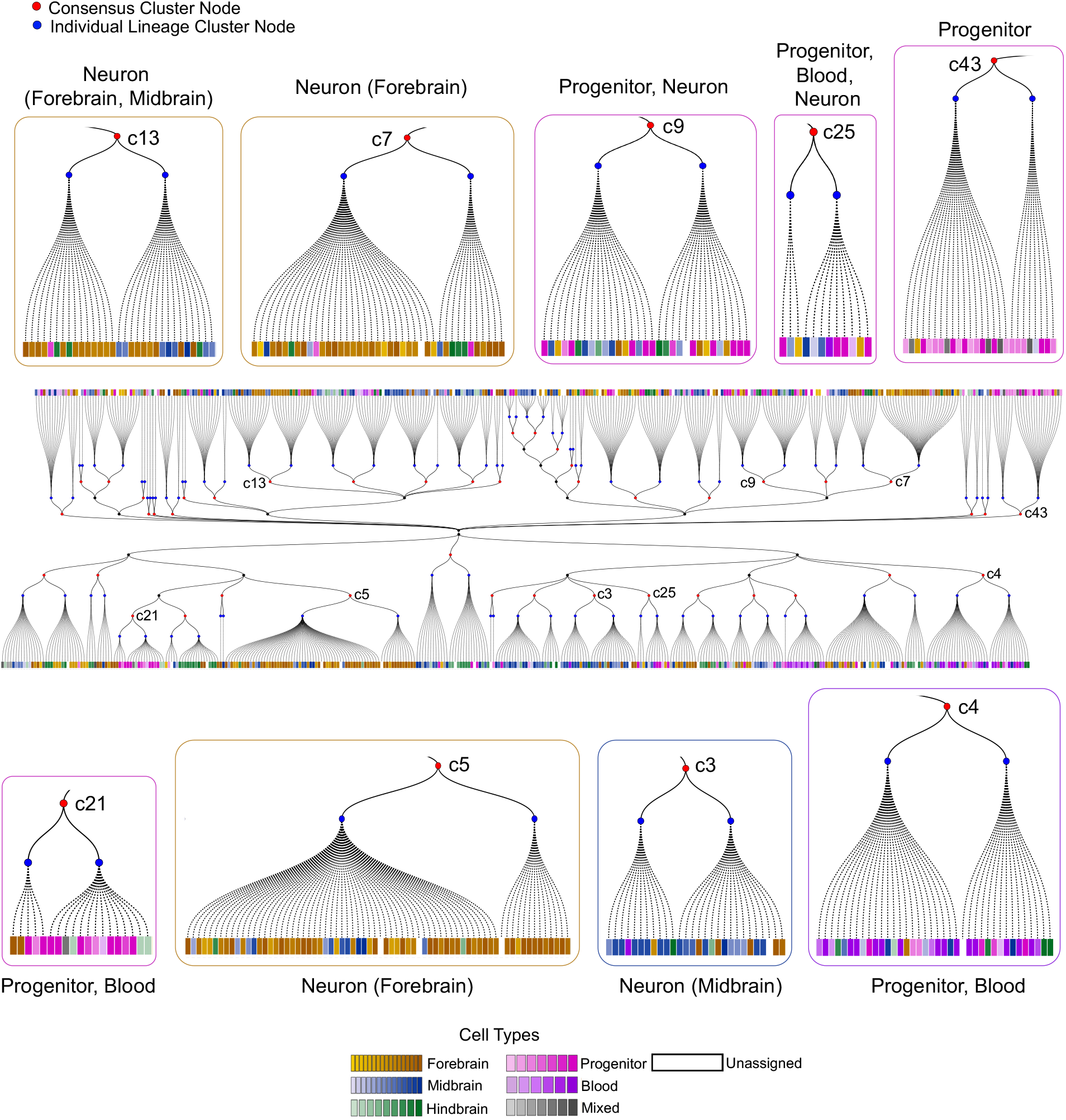
Consensus lineage tree for juvenile zebrafish brain. The two-sided tree in the middle represents the consensus tree generated by LinTIMaT by combining the individual trees for ZF1 and ZF3. Blue nodes here represent the clusters from individual fishes (left node: ZF1, right node: ZF3), and red nodes represent the matched consensus clusters. Each leaf node is a cell, represented by a square, and its color represents its cell type as indicated in the legend. Subtrees illustrate examples of successfully matched clusters from two individual lineage trees.

We first evaluated the consensus lineage by computing its Adjusted Rand Index (ARI) based on the 63 cell types obtained by (20). Our analysis showed that, similar to what we observed for the individual trees, when learning the consensus tree, minimizing the objective function improved the ARI score of the matched clusters (Supplementary Fig. S4). The individual fishes had different spatial distribution of cells (for example, ZF1 had more forebrain cells and ZF3 had more hindbrain cells) making it very difficult to achieve perfect cluster matching for all clusters. Despite this, the ARI for the consensus lineage (0.077) was comparable to the individual LinTIMaT lineages (0.084 and 0.076 for ZF1 and ZF3 respectively) and higher than both individual MP lineages (0.061 and 0.056 for ZF1 and ZF3 respectively). The consensus lineage preserved some of the ancestor-descendant relationship of the individual lineages (Supplementary Fig. S5) while in some cases it placed similar cell clusters from different branches of the individual trees under the same subtree (Supplementary Fig. S6). Thus, in addition to enabling the integration of data across experiments, by using more data, the consensus method can also help improve on the individual trees themselves.

We further analyzed the matched clusters for spatial enrichment. The clusters in the consensus lineage were enriched in all three regions of brain (hindbrain, forebrain and midbrain) as shown in Fig. 5a. The consensus lineage showed more enriched hindbrain clusters compared to that of ZF1 and more enriched forebrain clusters compared to that of ZF3. Some of the consensus clusters contained cells from two regions, for example, forebrain and midbrain. This likely occurred due to the uneven distribution of cells in different brain regions for the individual fishes.

**Figure 5:**
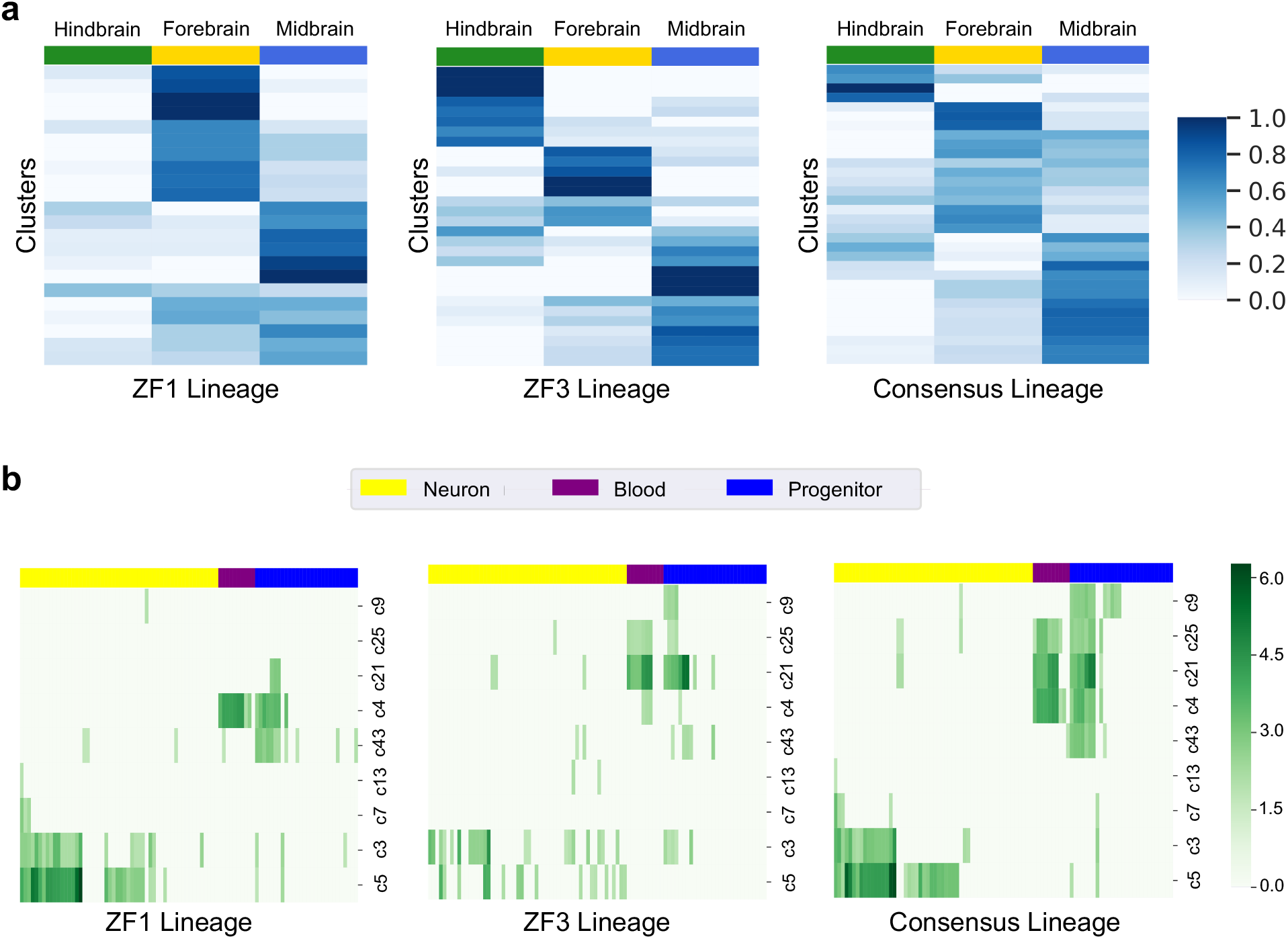
Functional analysis of cell clusters. (a) Heat map of the distribution of cell clusters for each region of the brain (columns). Cell types were classified as belonging to the forebrain, midbrain or hindbrain, and the proportions of cells within each region were calculated for each cluster. Each row sums to 1. Region proportions were colored as shown in key. The leftmost panel shows the heat map for the clusters in ZF1 lineage (subsampled), middle panel shows the heat map for ZF3 lineage and the rightmost panel shows the heat map for the consensus lineage. (b) Heat map of the p-values 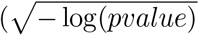, higher value means more significant) for GO terms for selected consensus clusters. Rows represent selected consensus clusters and columns represent different GO terms (Supplementary Table S4). Yellow, purple and blue columns correspond to GO terms related to neurons, blood and progenitors respectively. The leftmost panel shows the heat map for ZF1, middle panel for ZF3 and the rightmost panel for the consensus tree. As can be seen, the consensus tree correctly combines the unique terms identified for each tree. On one hand, it is able to identify neuron clusters, which are well represented in ZF1 but not in ZF3. On the other hand, it is able to identify progenitor clusters which are not well represented in ZF3.

To determine the biological significance of the consensus and individual lineage trees, we performed Gene Ontology (GO) analysis (Methods) on matched clusters that contained more than 10 cells. We also filtered the matched clusters where the individual cluster contained less than 3 cells. We selected all GO terms related to the three major cell types (neuron, blood and progenitor) present in the data (see Supplementary Table S3 and Supplementary Table S4 for the keywords and list of GO terms). Fig. 5b and Supplementary Fig. S7 presents the enrichment of the GO terms in the clusters in terms of p-values. The consensus clusters show coherent enrichment of GO terms for all three major cell types. For example, clusters c3 (midbrain), c5 (forebrain), c7 (forebrain) and c13 (forebrain and midbrain) had high p-value for the GO terms related to neuron but very low p-value for GO terms related to blood and progenitor. Similarly, cluster c43 displayed more enrichment of the progenitor GO terms. Clusters c4, c21 and c25 that consisted mostly of blood and progenitor cells showed enrichment of GO terms related to these two cell types. Cluster c9 consisted of mostly progenitor cells and some midbrain neurons, consequently it showed enrichment of mostly progenitor GO terms and a few neuron related GO terms. The coherence of enrichment can also be observed in the proportion of the GO terms related to the three major cell types (Supplementary Fig. S8). Clusters in the individual lineage trees also showed enrichment of the three cell types. However, as expected, the consensus lineage clusters uncovered more GO terms with more significant p-values compared to the individual lineage clusters.

## Discussion

Recent studies (20; 21; 22) combine two complementary technologies, CRISPR-Cas9 genome editing and scRNA-seq for elucidating developmental lineages at whole organism level. These experimental techniques rely on introducing random heritable mutations during cell division using CRISPR-Cas9 and lineage trees are reconstructed based on these mutations using traditional phylogenetic algorithms (24) on profiled cells.

While this exciting new direction to address a decades old problem in-vivo has already led to several interesting insights into organ development in multicellular organisms, it suffers from a number of challenges that make it difficult to accurately reconstruct lineages and to combine trees reconstructed from repeat experiments. First, the tree reconstruction is performed solely based on recovered mutation data, which might be noisy. In addition, the space for the mutations is limited resulting in saturation restricting the ability to further subdivide cells at later stages. Finally, due to the random nature of these mutations, it is impossible to utilize them to reconstruct a consensus lineage tree by combining data from repeated experiments of the same species, in contrast to most phylogenetic studies (32). No computational method has been developed to address these challenges.

To address these issues, we developed a new statistical method, LinTIMaT, which directly incorporates expression data along with mutation information for reconstructing both, individual and consensus lineage trees. Our method defines a global likelihood function that combines both mutation agreement and expression coherence. As we show, by optimizing this likelihood, LinTIMaT reconstructs lineage trees that are as good as the best mutation-only lineages while greatly improves over mutation-only lineages in terms of expression coherence, clade homogeneity and functional annotations. In addition, by employing agreement based on expression data, we further reconstruct a consensus lineage that retains most of the original tree branching for each individual while improving on the individual lineages by uncovering more biologically significant GO annotations corresponding to different major cell types.

Even though ground truth for developmental lineages is missing, we have validated the accuracy of the resulting trees using complementary information (global clustering based on six individuals and functional enrichment analysis). Our analysis shows that gene expression data can be very useful for selecting between several lineages with equivalent explanation of the mutation data. Since traditional phylogenetic maximum parsimony algorithms (24) as used in the original study (20) end up selecting a solution that is only slightly better or equivalent compared to several competing ones (though can be very different), the ability to use additional information (in our case gene expression) to select between these equally likely lineage trees is a major advantage of LinTIMaT. LinTIMaT’s Bayesian hierarchical model for gene expression data also provides a statistical method for inferring cell clusters with coherent cell types from the lineage tree. While it is not clear yet if all organisms follow the same detailed developmental plan as *c. elegans* (33), the ability to combine lineage trees studied in multiple individuals of the same species can lead to more general trees that capture the major branching events for the species. In addition, consensus trees can be used to improve branchings in the individual trees by combining information from multiple experiments. To the best of our knowledge, LinTIMaT’s solution, which is based on iteratively matching cell clusters based on their expression, is the first to enable the reconstruction of such consensus lineage trees from experiments that simultaneously profile lineage recordings and single-cell transcriptomes.

The application of LinTIMaT to zebrafish brain development illustrates its potential in delineating lineage relationships in complex tissues. The method is general and can work with data for any species for any tissue. While the joint profiling of lineage recordings and single-cell transcriptomes by experimental methods such as scGESTALT (20) laid the foundation for generating data suitable for identifying cellular relationships during development and disease, LinTIMaT provides the seminal computational approach for utilizing such data for accurate lineage reconstruction. As the usage of the experimental methods expands from zebrafish to other model organisms and human organoid samples (3), LinTIMaT would serve as a powerful component in the biologists’ toolbox in reconstructing more accurate and detailed lineages for investigating normal as well as pathological development.

## Methods

### Processing of the input data

LinTIMaT is designed for single-cell datasets in which both edited barcode and scRNA-seq data are available from the same cell. Each CRISPR-Cas9 mutation event (edit) has variable length and a single event could span across multiple adjacent sites. To construct a lineage tree from the mutation data we first count the number of unique synthetic markers (Cas9 edits) that occur in the 9 mutation sites. For each cell, the mutated barcode is represented by a binary vector of length equal to the number of unique synthetic markers, where each bit represents the state of a synthetic marker. For example, for Fish 1 in the scGESTALT dataset there are 324 entries in this vector for each cell. We use the mutation data to construct a paired-event matrix, *ε*_*B×S*_ for *B* unique barcodes and *S* unique editing events (synthetic markers), and an imputed gene-expression matrix, 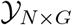 for *N* cells and *G* genes.

Each row of the paired-event matrix *ε*, corresponds to a mutated barcode (or allele) and each column corresponds to a unique editing event. An entry *e*_*bs*_ of *ε* is a binary variable that denotes the presence or absence of marker *s* in barcode *b* (1 or 0). Each cell *c* is associated with one, and only one, of the *B* unique barcodes. As a result, each barcode represents a group of cells. For each cell *c* = 1,…, *N*, *z*_*c*_ denotes the barcode *b* profied for that the cell, *z*_*c*_ = *b*, where *b* ∈ {1,…, *B*}. Thus, the matrix *ε* can be transformed to an *N* × *S* matrix for *N* cells and *S* markers, where the row *c* will correspond to the barcode *z*_*c*_ associated with cell *c*.

The other type of data our method uses is scRNA-seq data. In general, the method can work with any such data. For the specific data used in this paper, we observed a high dropout rate (94% entries were 0). To address this issue we tested a number of imputation methods (see Supplementary Methods and Supplementary Fig. S9) and selected DrImpute (34) for imputation. DrImpute first clusters the data, and then each zero expression value is imputed with the mean gene expression of the cells in the cluster the cell belongs to. Next, we normalized the expression of each cell and log2-transformed the results (Supplementary Methods).

### Likelihood of a cell lineage tree

As mentioned in the Introduction, our method aims to reconstruct a cell lineage tree by combining two complementary types of data. For this, we defined a joint likelihood function for the two data types and then search the space of possible trees for a model that maximizes the likelihood function. We first describe the likelihood function for each of the data types and then discuss how to perform a search for maximizing the joint likelihood to reconstruct the most likely tree.

### Cell lineage tree

We assume that the cell lineage tree is a rooted directed tree 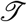. The root of this lineage tree denotes the initial cell that does not contain any marker (or editing event). The leaves of this tree denote cells profiled in the experiments. Cells go through the differentiation process along the branches of the lineage tree and as part of this process acquire the synthetic mutations (edits) induced by Cas9. Some of the internal nodes in the cell lineage tree represent the unique mutated barcodes shared by the leaves (cells) under that specific internal node. For ease of computation, we first reconstruct a rooted binary lineage tree and later eliminate the internal branchings that are not supported by any synthetic mutations.

### Mutation likelihood

The first component of the likelihood function evaluates the likelihood of the cell lineage tree based on the mutation data. The mutations induced by Cas9 are irreversible since the Cas9 protein cannot bind to the target sites once changed. To account for this, we impose a Camin-Sokal parsimony criterion (35) on each synthetic mutation. This criterion states that each synthetic mutation can be acquired at least once along the lineage but once acquired they are never lost. We also assume that the synthetic mutations are acquired independently and parsimoniously as higher number of mutations along the branches of the cell lineage indicates a more complex mutational history which is less likely. For a given cell lineage tree 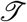, we first use Fitch’s algorithm (36) to assign ancestral states for each marker to each internal node of the tree satisfying maximum parsimony. Such an assignment, 𝓐 results in the least number of mutations on the given tree. The mutation likelihood (𝓛_*M*_) of the cell lineage tree is then given by

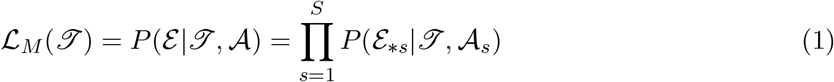

where *ε*_∗*s*_ is the observed data for marker *s* which is a vector corresponding to *N* values for *N* cells. 𝓐_*s*_ denotes the parsimonious assignment of ancestral states for all internal nodes for marker *s*.

For an internal node *v* with children *u* and *w*, 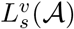 denotes the partial conditional likelihood for marker *s* defined by

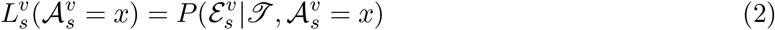

where 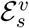 denotes the restriction of observed data for marker *s*, *ε*_∗*s*_ to the descendants of node *v* subject to the condition that 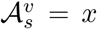 is the ancestral state for marker *s* assigned by Fitch’s algorithm, *x* ∈ {0, 1}. 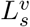 gives the likelihood for marker *s* for the subtree rooted at node *v*, given the assignment of ancestral states by Fitch’s algorithm.

The likelihood for the full observed data *ε*_∗*s*_ for marker *s* is given by

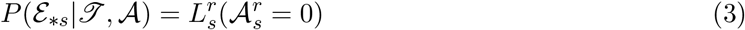

where *r* is the root of the lineage tree. Since, the root of the tree does not contain any synthetic mutation, 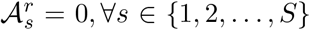. For any internal node *v* with children *u* and *w*, the partial conditional likelihood satisfies the recursive relation

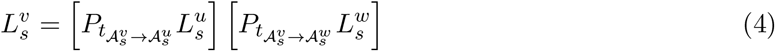

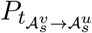 and 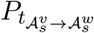 denote the transition probabilities on branches that connect *v* and *u*, and *v*and *w* respectively. For each synthetic mutation *s*, we define a transition probability matrix given by

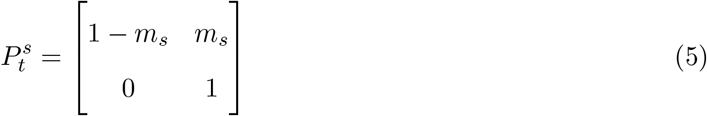

where *m*_*s*_ denotes the fraction of cells harboring *s* and 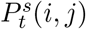 denotes the probability of transition from state *i* to state *j* along any branch of the tree. If a mutation assignment violates the Camin-Sokal parsimony criterion (i.e. a mutation is reversed), the log-likelihood is heavily penalized (−100000) so that LinTIMaT prefers the tree without such violation.

For each leaf *l* of the tree, the partial likelihood is set to 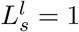.

### Expression likelihood

For the expression data likelihood, we model the lineage as a Bayesian hierarchical clustering (BHC) (25) of the cells and used the likelihood formulation provided by BHC. BHC is a bottom-up agglomerative clustering method that iteratively merges clusters based on marginal likelihoods. Following several other methods we assume a diagonal matrix when computing gene expression variance for each internal and leaf node (37; 38). Following BHC algorithm, we compute the marginal likelihoods of all the partitions consistent with the given lineage tree based on a Dirichlet process mixture model. The expression likelihood for a particular gene is given by the marginal likelihood for the root of the tree and it essentially provides a lower bound on the marginal likelihood of a Dirichlet process mixture model. The product of the likelihoods for all the genes is used to determine the expression likelihood (*𝓛*_*E*_) for the complete dataset.

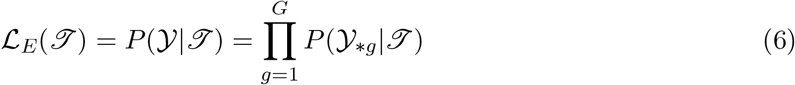

where 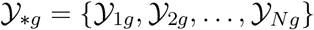 is the vector containing expression values for gene *g* for all cells. 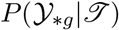 is the expression likelihood for the lineage tree which is also the marginal likelihood 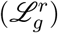 for the root of the tree

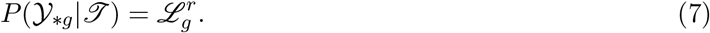

For an internal node *v* with children *u* and *w*, 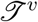 denotes the subtree rooted at *v*. Let 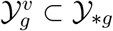 be the set of gene expression data at the leaves under the subtree 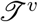 and 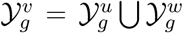. To compute the marginal likelihood for node *v* 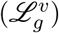, we compute the probability of the data under two hypotheses of BHC. The first hypothesis, 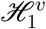 assumes that each data point is independently generated from a mixture model and each cluster corresponds to a distribution component. This means that the data points **y**^(*i*)^ in the cluster 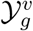 are independently and identically generated from a probabilistic model *P* (**y**|*θ*) with parameters *θ*. Thus, the marginal probability of the data 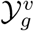 under the hypothesis 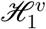 is given by

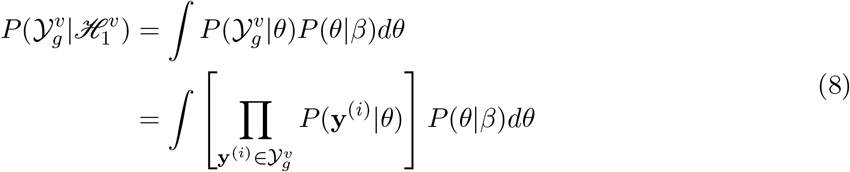

The integral in Eq. (8) can be made tractable by choosing a distribution with conjugate prior, as discussed in Supplementary Methods.

The alternative hypothesis 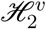 assumes that there are two or more clusters in 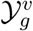. Instead of summing over all (exponential) possible ways of dividing 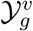 into two or more clusters, we follow the strategy in BHC (25) and sum over the clusterings that partition the data 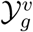 in a way that is consistent with the subtrees 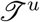 and 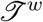. This gives us the probability of the data under the alternative hypothesis

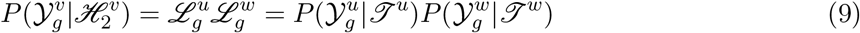

In Eq. (9), 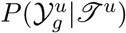 and 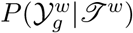 represents the marginal likelihoods of subtrees rooted in nodes *u* and *w* respectively. Combining the two likelihoods of the two hypotheses leads to a recursive definition of the marginal likelihood for the subtree 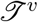 rooted at the node *v*

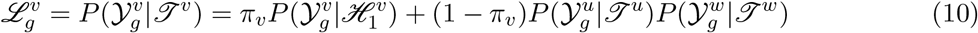

Where *π*_*v*_ is a parameter learned for weighting the two alternatives and is defined recursively for every node. The recursive definition of *π*_*v*_ for node *v* is given by

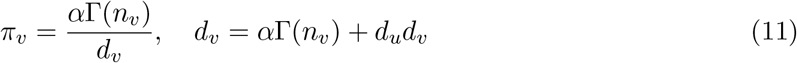

In Eq. (11), *α* denotes a hyperparameter, the concentration parameter of the Dirichlet process mixture model, *n*_*v*_ is the number of data points under the subtree 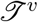 and Γ(.) is the Gamma function. For each leaf *l*, we set the values *π*_*l*_ = 1 and *d*_*l*_ = *α*. Also, for each leaf *l*, the marginal likelihood 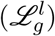 is calculated based on only the first hypothesis

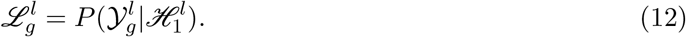

See Supplementary methods for discussion on how the prior is set for this model.

### Combined likelihood

For a given lineage tree, the joint log-likelihood (*𝓛*_*T*_) function for the mutation and expression data is a weighted sum given by

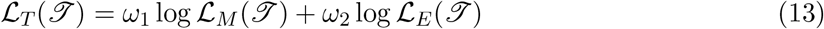

The values of *ω*_1_ and *ω*_2_ are chosen so that the values of the two likelihood components stay in the same range. In our experiments, we have used *ω*_1_ = 50 and *ω*_2_ = 1 (see Supplementary Fig. S10).

### Search algorithm for inferring lineage tree

Searching for the optimal tree under a maximum-likelihood framework like ours is a NP hard problem (39). We have thus developed a heuristic search algorithm which stochastically explores the space of lineage trees. The search algorithm consists of several stages as described below.

1. In the first step, we only focus on the barcodes and search for top scoring solutions. The search process starts from a random tree topology built on *B* leaves corresponding to *B* unique barcodes. In searching the barcode lineage tree, we employ the mutation likelihood function. In each iteration, a new barcode lineage tree, 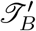 is proposed from the current tree 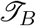 as we discuss below. If the proposed tree results in a higher likelihood, it is accepted, otherwise rejected. Instead of storing a single solution, we keep several of top scoring barcode lineage trees.

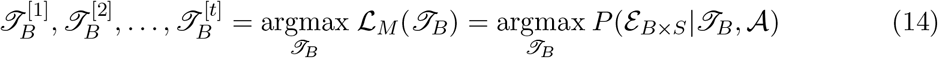
2. Next, we utilize the expression data. As mentioned above, a barcode can be shared between multiple cells. We thus next search for the best cellular subtree 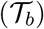 for the set of cells associated with each mutated barcode *b*. We employ hill-climbing to obtain single solution for each barcode that harbors more than 2 cells.

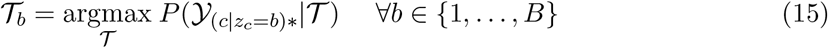
3. In the third step, we construct complete cell lineage trees by attaching cellular subtrees for each barcode to barcode lineage trees. To obtain the cell lineage tree 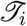 from a barcode lineage tree 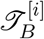, for each barcode *b* harboring more than 2 cells, we choose the cellular subtree 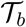 inferred in step 2 and connect its root to the leaf in 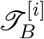 that corresponds to *b*. For a barcode *b* shared by two cells, the cells are connected to the leaf representing *b* in 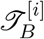 as children. This gives us *t* full binary cell lineage trees corresponding to *t* barcode lineage trees. Next, we evaluate the total log-likelihood of each of these cell lineage trees and choose the best one.

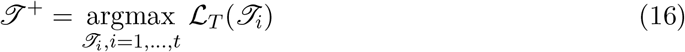 We also record the best mutation log-likelihood, 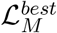 for the best cell lineage tree and define a threshold value for mutation log-likelihood

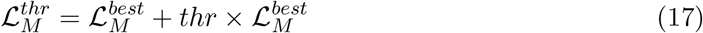

where, *thr* is a user-defined value close to 0.
4. In the final step, we perform another hill-climbing search to optimize the cell lineage tree 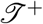 inferred in step 3 in terms of the joint likelihood function. The search starts from 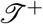 and in each iteration, we propose a new cell lineage tree 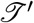 from the current tree 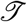 as we discuss below. For the new tree, we first ensure that the mutation log-likelihood of the new tree does not go below 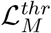. If this condition is satisfied and the total likelihood is improved then the new lineage tree is accepted. We stop the search if the total likelihood does not improve for a large number of iterations and return the best lineage tree achieved so far.

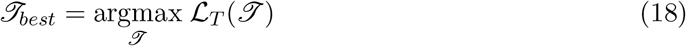

### Tree search moves

To explore the space of lineage trees, LinTIMaT employ two different types of moves that can make small and big changes in the tree topology. For this, we adopt two of the tree proposals described in (40) for efficient exploration of tree space for Bayesian phylogenetic inference. Both of these moves are branch-rearrangement proposals that alter the topology of the lineage tree.

The first tree proposal is a swapping move called Stochastic Nearest Neighbor Interchange (stNNI). In this move, we choose an internal branch as the focal branch and stochastically swap the subtrees attached to the focal branch. This type of move results in minimal topology change and is used only in the second step of our algorithm that infers cellular subtree for each mutated barcode.

The second tree proposal is a pruning-regrafting move, namely Random Subtree Pruning and Regrafting (rSPR). In this move, we first randomly select an interior branch, prune a subtree attached to that branch, and then reattach the subtree to another regrafting branch present in the other subtree. The regrafting branch is also chosen randomly. This type of move can introduce a larger amount of topology change in the tree and this is used in step 1 and 4 of our search algorithm.

### Inferring clusters from cell lineage tree

To obtain cell clusters from the inferred lineage tree, we employ the statistical model comparison criterion provided by the BHC model for gene expression data. For an internal node *v* with children *u* and *w*, we compute the probability of the data under two hypotheses. The first hypothesis suggests that all the cells under the node *v* belongs to a single cluster. We compute the posterior probability (*r*_*v*_) of this hypothesis using Bayes rule:

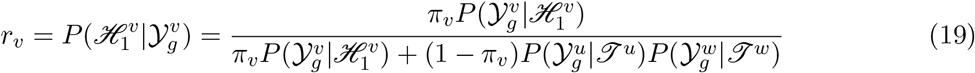

The lineage tree can be cut at the nodes where *r*_*v*_ goes from *r*_*v*_ < 0.5 to *r*_*v*_ > 0.5 to obtain clustering of cells.

### Combining lineage trees from multiple individuals to reconstruct a consensus lineage tree

As mentioned in the Introduction, a key challenge when working with CRISPR mutation data is the fact that these are not the same across different experiments. Thus, standard phylogenetic consensus tree building cannot be applied to this data. Instead, given a set of lineage trees, 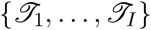 for *I* individuals, we construct a single lineage tree 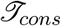 that jointly explains the differentiation of these individual organisms. Individual lineage trees that are input to the consensus lineage reconstruction method are built on a leaf set of different number of cells. 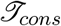 is constructed by following the steps below.

1. For each individual lineage tree 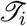, we infer the cell clusters based on gene expression data. Let us assume that *C*_*i*_ is the number of clusters inferred from lineage tree 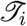. We define *K* as 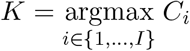.
2. For each individual lineage tree 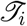 for which *C*_*i*_ < *K*, we split clusters to obtain *K* clusters. Splitting is done in decreasing order of the posterior probability *r*_*v*_ until the desired number of clusters is reached.
3. For each individual lineage tree 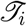, we obtain the backbone tree 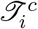 built using these *K* clusters.
4. 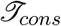 is a lineage tree built on a leaf set of *K* clusters. We first define a cluster matching 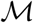 as a matching where each cluster in each individual lineage tree 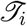 (or each leaf in 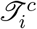) is matched with a leaf of 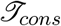. We reconstruct 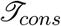 and a cluster matching 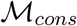 by minimizing an objective function given by

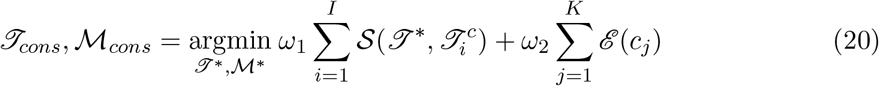

where 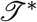 is a candidate consensus lineage, 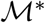 is a candidate cluster matching, 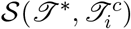 denotes the sum of pairwise leaf shortest path distance between candidate consensus lineage 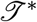 and individual lineage 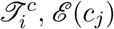 denotes the sum of pairwise distance between the clusters of the individual lineage trees that match with cluster (or leaf) *c*_*j*_ in the candidate consensus lineage. The objective function for searching the consensus lineage and the optimal cluster matching is described below in detail. We employ a two-step heuristic search algorithm for optimizing the objective function (described below).

### Objective function for searching consensus lineage tree

The objective function for reconstructing the consensus lineage attempts to balance two competing issues. The first is that the consensus tree should be as close as possible to each of the individual lineages. The second is that the agreement (in terms of expression) between nearby subtrees in the consensus tree would be high. We thus attempt to minimize two different distance functions to select the optimal tree. 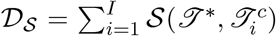 computes the distance (or disagreement) between *i* the topology of the consensus lineage and the individual lineage trees. 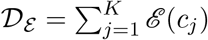 is the other distance function which attempts to minimize disagreement between the gene expression values of matched clusters.

For computing *𝓓*_*S*_, we employ the sum of pairwise leaf shortest-path distance (41; 42) between two trees as a distance measure for comparing two tree topologies. The shortest path distance *δ*_*ij*_(.) between two leaves *c*_*i*_ and *c*_*j*_ in a tree is given by the sum of the number of edges that separate them from their most recent common ancestor. Overall pairwise leaf shortest-path distance between two trees is obtained by summing up the absolute differences between the shortest-path distances of all unordered pairs of leaves in the two trees

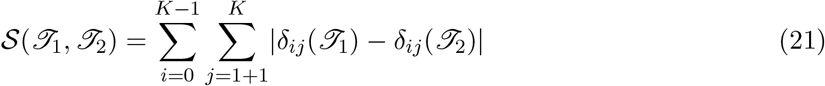

For computing *𝓓*_*ε*_, we sum the pairwise distance between the clusters of the individual lineage trees that match with a leaf of the consensus lineage. 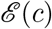 is given by

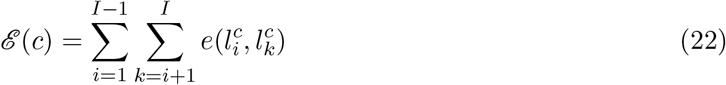

where 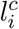 and 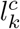 denote clusters in individual lineages that match with leaf *c* in candidate consensus lineage. *e*(.) denotes the Euclidean distance between the gene expression value of two clusters.

### Search algorithm for inferring consensus lineage

We use a two-step heuristic search algorithm for inferring the consensus lineage and the corresponding cluster matching.

1. The first step employs an iterative search. In each iteration, we first find a better cluster matching (see Supplementary Methods for details) than the current matching 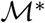, and then keeping this matching fixed, we improve the topology of the consensus tree. It is important to note that, a new cluster matching modifies both *𝓓*_*ε*_ and 𝓓_*S*_, whereas a new tree topology modifies only 𝓓_*S*_. This iterative search goes on until cluster matching can not be improved further. Let us assume, 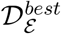 is the distance corresponding to the best cluster matching achieved. We define a threshold value for the cluster matching distance

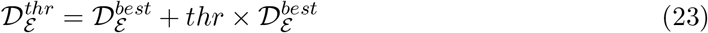
2. In the second step, we try to improve the consensus lineage by improving the objective function 𝓓_*S*_ + 𝓓_*ε*_ using a stochastic search. In the joint 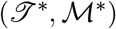 space, we consider two types of moves to propose a new configuration. In each iteration, from the current configuration 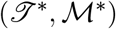, we either propose a new matching (Supplementary Methods) 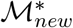 or a new tree topology 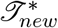 using the tree search moves. When a new matching 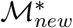 is proposed, we first ensure that the cluster matching distance for the new matching does not lead to values above the threshold 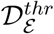. If this condition is satisfied and the objective function is minimized then the new matching is accepted. If the proposed tree topology 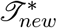 achieves lower value for the objective function, it is accepted. The search procedure terminates when the objective function does not improve or the maximum number of iterations has been reached.

### GO analysis on clusters identified by LinTIMaT

To perform GO (Gene Ontology) analysis on consensus lineage clusters, we first identify a set of differentially expressed (DE) genes based on t-test of 2 groups of cells. The first group consists of the cells in the consensus cluster and the second group includes all other cells in the dataset. From the set of DE genes, we further select the genes that have higher mean expression in the first group, with a p-value smaller than 0.05 (or top 500 if more than 500 genes achieve this p-value). Finally, we use gprofiler (43) to perform GO query for the genes selected for each cluster.

### Analyzing the cell clustering performance of a lineage tree

For assessing the cell clustering performance of a lineage tree, we use 63 cell types obtained by (20) as ground truth and use Adjusted Rand Index (ARI) as the clustering metric following (44). Basically, ARI is calculated based on the number of agreements and number of disagreements of two groupings, with randomness taken into account. ARI is defined as follows. Let *X* = {*X*_1_, *X*_2_,…, *X*_r_}, *Y* = {*Y*_1_, *Y*_2_,…, *Y*_s_} be two groupings, where *X* has *r* clusters and *Y* has *s* clusters. We can set the overlap between *X* and *Y* using a table *N* with size *r* ∗ *s*, where *N*_*ij*_ = |*X*_*i*_ ∩ *Y*_*j*_| denotes the number of objects that are common to both *X*_*i*_ and *Y*_*j*_. Let *a*_*i*_ = ∑_*j*_ *N*_*ij*_, *b*_*j*_ = ∑ _*i*_ *N*_*ij*_, *n* be the total number of samples, then ARI is given by

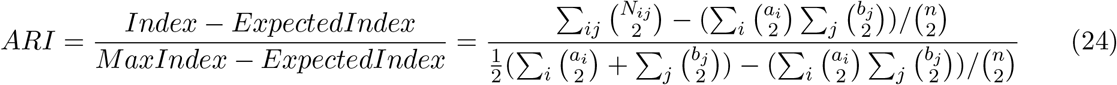

### Software availability

LinTIMaT has been implemented in Java and is freely available at https://github.com/jessica1338/LinTIMaT, under the MIT license. This implementation uses the PhyloNet (45) library.

### Data availability

The high-throughput datasets used in this study were previously deposited in the Gene Expression Omnibus under accession number GSE105010. Lineage trees are available for exploring at https://jessica1338.github.io/LinTIMaT/

## Acknowledgements

This work was partially funded by the National Institutes of Health (NIH) [grants 1R01GM122096 and OT2OD026682 to Z.B.J.]. The order of authorship of the first two authors was determined by a coin flip.

