## Supplementary Material for "Single-cell Lineage Tracing by Integrating CRISPR-Cas9 Mutations with Transcriptomic Data"

Hamim Zafar[1,†], Chieh Lin[2,†], Ziv Bar-Joseph[1,2,*]
[1]Computational Biology Department, [2]Machine Learning Department
School of Computer Science, Carnegie Mellon University
[†] *Authors contributed equally*

May 6, 2019

**3 Supplementary Figures** **6**

**4 Supplementary Tables** **16**

### 1 Supplementary Methods

#### 1.1 Imputation of gene expression data

The scRNA-seq data in the scGESTALT datasets displayed a high dropout rate (94% entries were 0). To address this, we decided to impute the scRNA-seq data. To ensure that LinTIMaT's likelihood function is not sensitive to the imputation method, we tested two different imputation methods named DrImpute [1] and SAVER [2]. We observed that LinTIMaT's results were not dependent on the imputation method used and the cell clustering performance from the same lineage tree were similar for the different imputation methods (Supplementary Fig. S9). We finally selected DrImpute as it has been shown to perform better in other experiments [3].

#### 1.2 Normalization of scRNA-seq data

After imputing the scRNA-seq data using DrImpute, we normalized the data. For normalization, we divided each expression value by the sum of expression values for each cell and multiplied the expression value with a scaling factor 10000, which is a common scaling factor for UMI (unique molecular identifier) data.

#### 1.3 Prior distribution for computing expression likelihood

LinTIMaT's expression likelihood function computes the probability of the expression data under a node based on two alternative hypotheses. The first hypothesis computes the marginal probability of the data being generated from a single cluster. For computing this marginal probability using Eq. (8), we choose a univariate Gaussian distribution and the Normal-inverse-chi-squared (NIX) prior for $\beta$. NIX prior is the univariate version of normal-inverse-Wishart (NIW) prior, which is the prior suggested by BHC [4]. NIX prior has the following parameters $\beta = (\mu_0, \kappa_0, \sigma_0^2, \nu_0)$, where $\mu_0$ and $\sigma_0^2$ are the priors on the mean and variance of the Gaussian distribution. $\kappa_0$ and $\nu_0$ are the confidence on the prior of the mean and variance respectively. The posterior parameters $\{\mu_v, \sigma_v^2, \kappa_v, \nu_v\}$ and the marginal probability for the subtree $\mathscr{T}^v$ rooted at node $v$ under the NIX prior are derived according to [5] and shown below

$$\kappa_v = \kappa_0 + n_v \tag{1}$$

$$\nu_v = \nu_0 + n_v \tag{2}$$

$$\mu_v = \frac{\kappa_0 \mu_0 + n_v \bar{\mathbf{y}}_v}{\kappa_v} \tag{3}$$

$$\sigma_v^2 = \frac{1}{\nu_v} \left( \nu_0 \sigma_0^2 + \sum_{\mathbf{y}^{(i)} \in \mathcal{Y}_g^v} (\mathbf{y}^{(i)} - \bar{\mathbf{y}}_v)^2 + \frac{n_v \kappa_0}{\kappa_v} (\mu_0 - \bar{\mathbf{y}}_v)^2 \right), \tag{4}$$

where $\bar{\mathbf{y}}_v$ is the sample mean of $\mathcal{Y}_g^v$. The marginal likelihood is given by

$$P(\mathcal{Y}_g^v | \mathscr{H}_1^v) = \frac{\Gamma(\nu_v/2)}{\Gamma(\nu_0/2)} \sqrt{\frac{\kappa_0}{\kappa_v}} \frac{(\nu_0 \sigma_0^2)^{\nu_0/2}}{(\nu_v \sigma_k^2)^{\nu_v/2}} \frac{1}{\pi^{n_v/2}} \tag{5}$$

For the hyperparameters of $\beta$, we set $\mu_0$ and $\sigma_0^2$ to sample mean and sample variance based on all single cells, $\mu_0 = \frac{1}{N} \sum_{c=1}^{N} \mathcal{Y}_{cg}, \sigma_0^2 = \sum_{c=1}^{N} (\mathcal{Y}_{cg} - \mu_0)^2$. The confidence parameters are set to 1, $\kappa_0 = \nu_0 = 1$.

##### 1.4 Proposal for cluster matching

We use a two-step heuristic search algorithm for inferring the consensus lineage and the corresponding cluster matching. We employ two different proposals for proposing a new cluster matching for the two steps of the consensus lineage search algorithm respectively.

1. Let us assume $\mathcal{M}_{old}$ denotes the current matching from which we want to propose a new cluster matching. In $\mathcal{M}_{old}$, $K$ clusters (leaves) in the consensus lineage are

matched to $K$ clusters in each individual lineage. Let $\{c_1, \ldots, c_K\}$ denotes the $K$ clusters in the consensus lineage. In $\mathcal{M}_{old}$, for individual lineage $\mathscr{T}_i^c$ ($i \in \{1, \ldots, I\}$), let $l_i^{c_x}$ and $l_i^{c_y}$ denote the clusters that match with clusters $c_x$ and $c_y$ in the consensus respectively. We can propose a new cluster matching $\mathcal{M}_{new}^i$ by swapping the matchings of $l_i^{c_x}$ and $l_i^{c_y}$ with $c_x$ and $c_y$. There are $\mathcal{O}(K^2)$ such possible swaps. For each such swap $\eta$, we compute a score $\Sigma_\eta = \Delta_\mathcal{S}^\eta + \Delta_\mathcal{E}^\eta$, where $\Delta_\mathcal{S}^\eta$ denotes the improvement in $\mathcal{D}_\mathcal{S}$ after the swap and $\Delta_\mathcal{E}^\eta$ denotes the improvement in $\mathcal{D}_\mathcal{E}$ after the swap. The swaps for which both $\Delta_\mathcal{E}^\eta$ and $\Sigma_\eta$ are positive are considered to be good swaps. One such good swap is chosen randomly to propose a new matching $\mathcal{M}_{new}^i$. If no good swap is available, no swapping is performed. This is done sequentially for all $I$ individual lineages to produce a new cluster matching $\mathcal{M}_{new}$.

2. In the second step of the search algorithm, we perform a random swap to propose a new matching. For consensus lineage, we randomly choose two clusters $c_x$ and $c_y$, and in the individual lineage $\mathscr{T}_i^c$, we swap their matchings with $l_i^{c_x}$ and $l_i^{c_y}$.

##### 1.5 Visualizing LinTIMaT trees

Following [6], individual cells (leaves) in the lineage trees were annotated by their corresponding cell types. LinTIMaT lineage trees were converted into JSON objects with annotated cell type membership using custom python scripts. Finally, the JSON objects were visualized using the modified custom scripts of [6] using D3 software framework. The visualization web page also displays additional information on each tree node such as mutations and cell type proportions.

### 2 Supplementary Results

#### 2.1 Computing ARI for cell clustering based on Maximum Parsimony lineage trees

To compare LinTIMaT lineages against Maximum Parsimony (MP) lineages from [6] for ZF1 and ZF3, we compared the cell clustering performance for the lineage trees. The cell clustering performance was measured by computing ARI. While LinTIMaT trees allow for inferring cell clusters based on gene expression data, MP lineage trees do not provide such option. For MP lineage trees, the unique barcodes can be treated as cell clusters. However, to be more thorough, we also cut the MP trees at different levels to obtain different possible cell clusterings. The ARI values for the clusterings obtained by cutting the MP trees at level 1-6 and the barcode level for both ZF1 and ZF3 are shown in Supplementary Table S1. For both fishes, the barcode level clustering achieved the highest ARI values. Consequently, these values were used for comparing MP trees against LinTIMaT trees.

#### 2.2 ARI values for lineages reconstructed by LinTIMaT

We computed Adjusted Rand Index (ARI) for the cell clustering inferred from lineages reconstructed using LinTIMaT. ARI was 0.084 for ZF1 and 0.076 for ZF3 respectively.

#### 3 Supplementary Figures

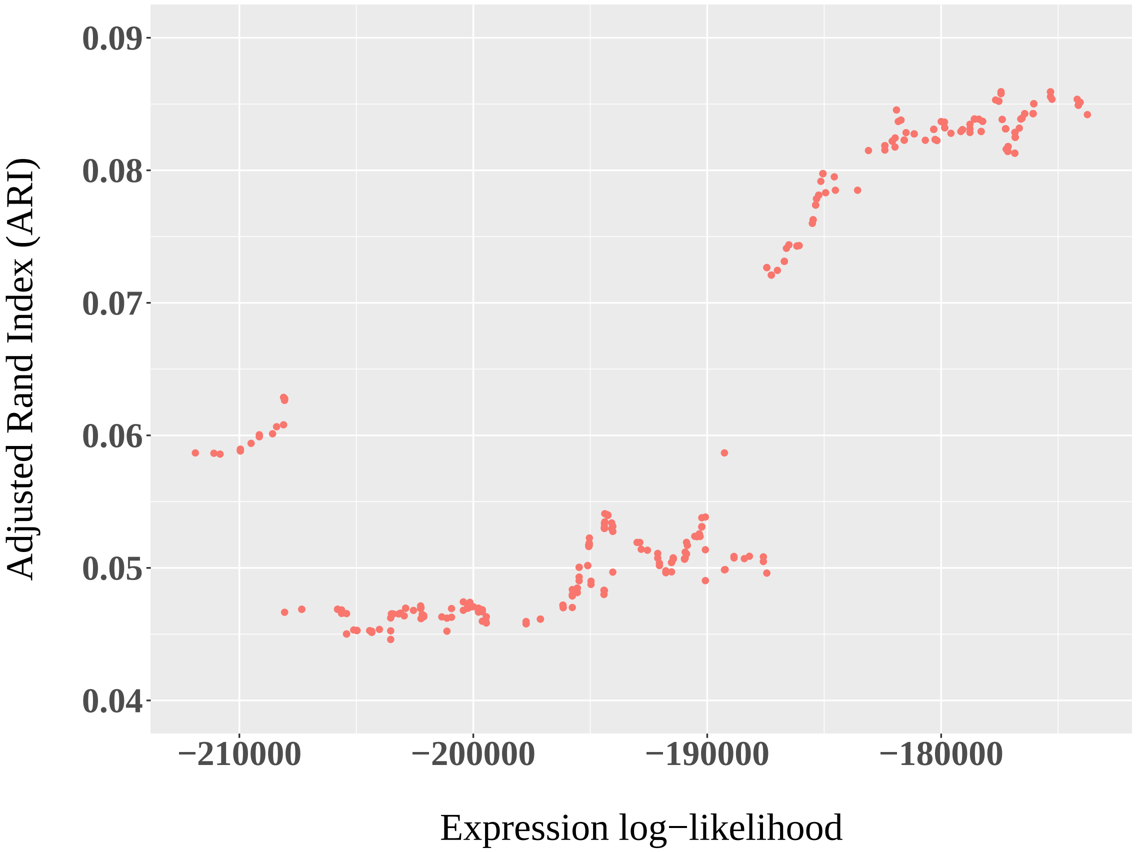

Supplementary Figure S1: Adjusted Rand Index (ARI) as a function of the expression likelihood score calculated by LinTIMaT for ZF1.

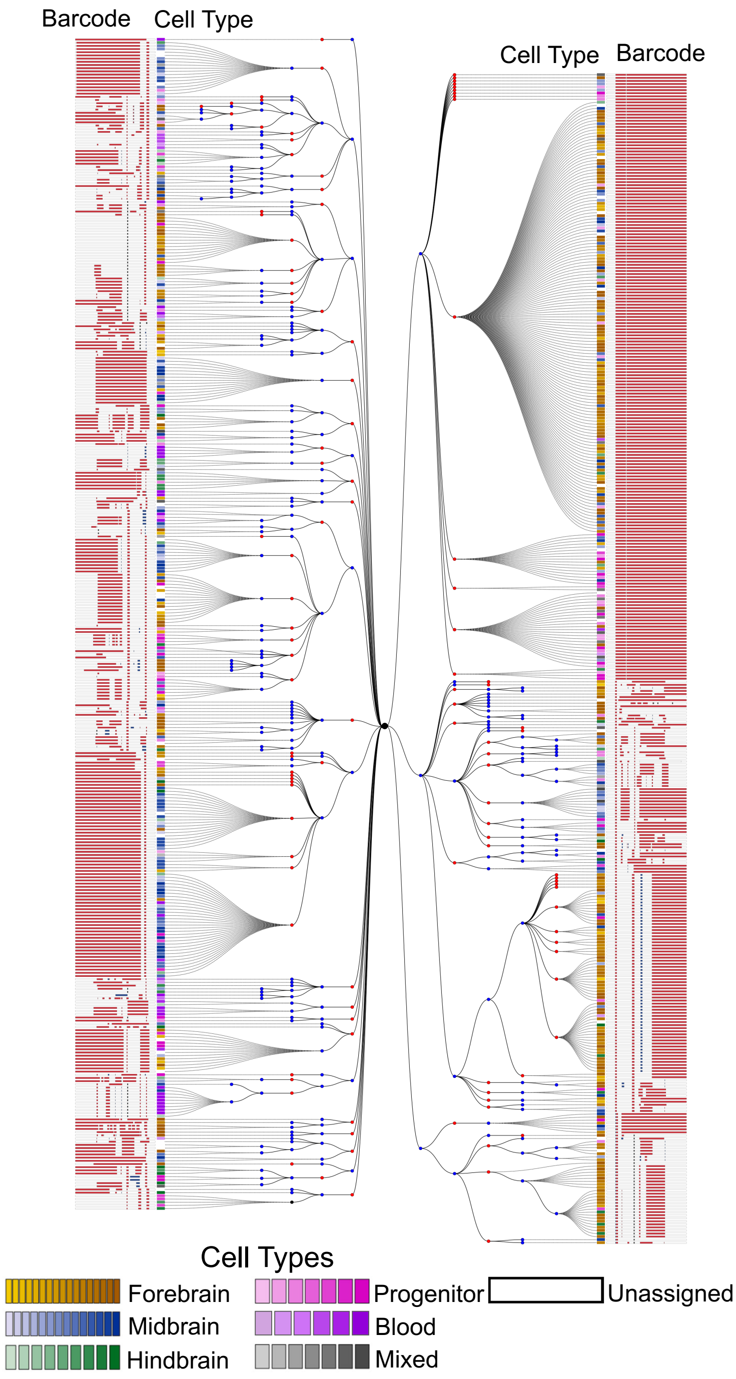

Supplementary Figure S2: The lineage tree reconstructed by LinTIMaT for a single juvenile zebrafish brain (ZF1). The lineage tree is built on 750 cells. Blue nodes represent Cas9-editing events (mutations) and red nodes represent clusters inferred by LinTIMaT from transcriptomic data. Each leaf node is a cell, represented by a square, and its color represents its cell type as indicated in the legend. The mutated barcode for each cell is displayed as a white bar with insertions (blue) and deletions (red).

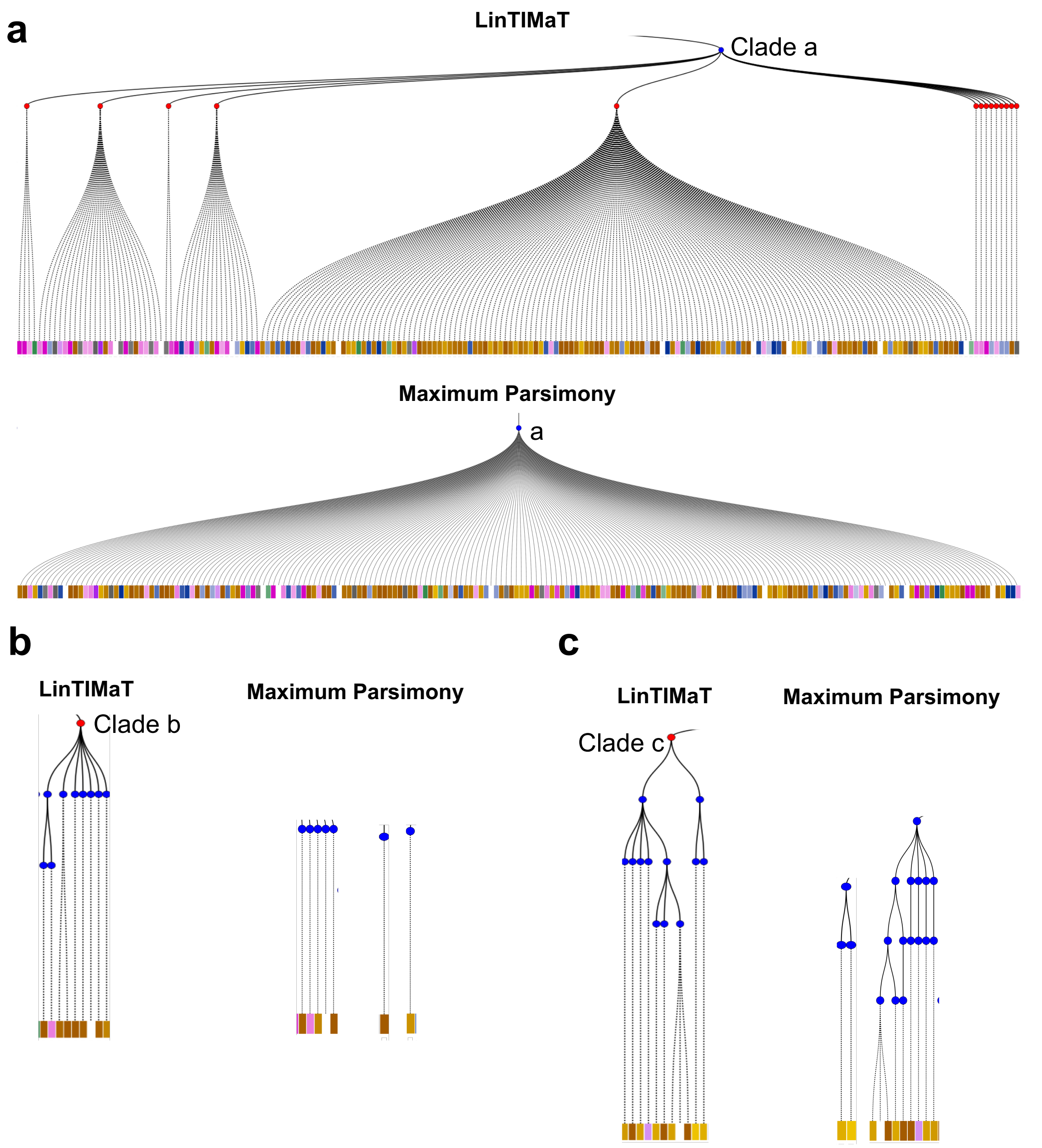

Supplementary Figure S3: Example subtrees in the lineage tree reconstructed by LinTIMaT for a single juvenile zebrafish brain (ZF1). (a) Example subtree shows that by using transcriptomics data LinTIMaT is able to further refine subtrees in which all cells share the same barcode which can help overcome saturation issues. (b-c) Example subtrees displaying LinTIMaT's ability to cluster cells with different barcodes together based on their cell types, maximum parsimony puts them on distinct branches.

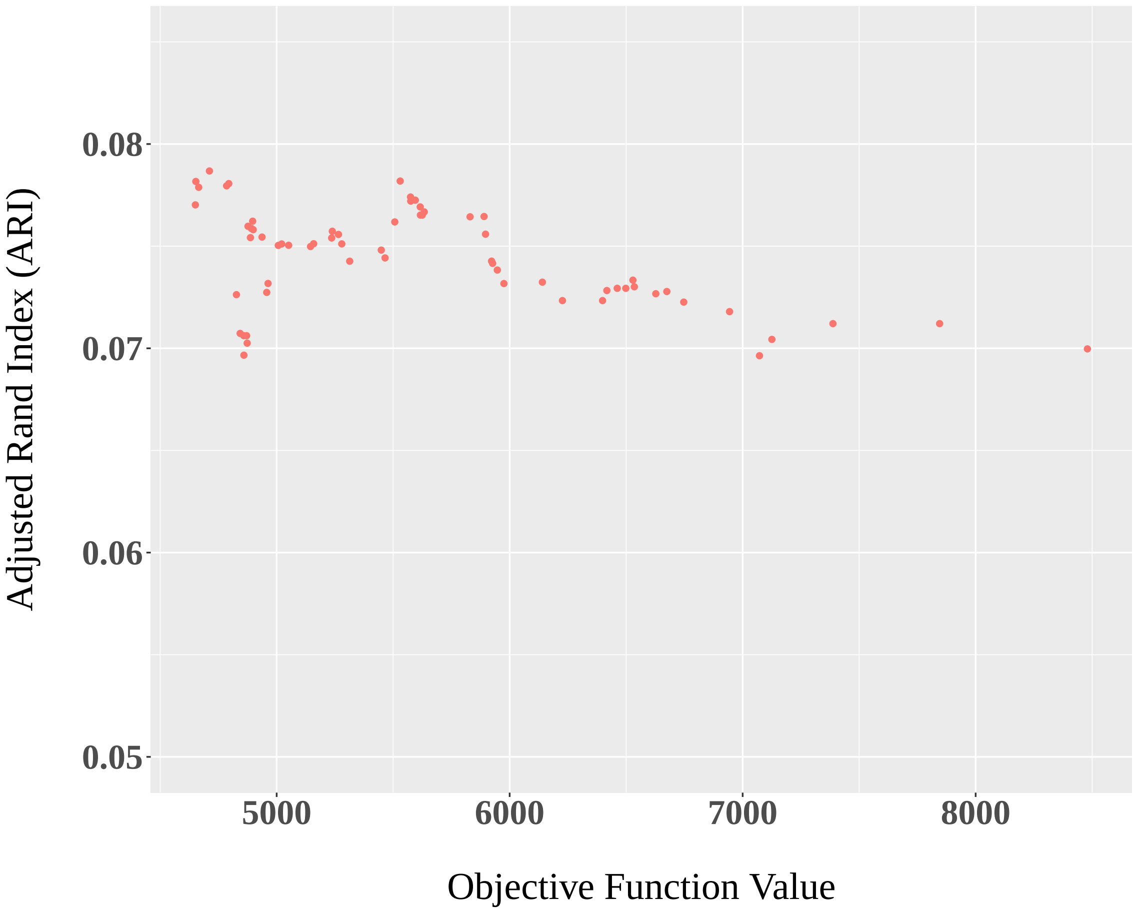

Supplementary Figure S4: Adjusted Rand Index (ARI) as a function of the value of the objective function for reconstructing consensus lineage. Lower value of the objective function improves ARI and leads to better cluster matching. The inferred consensus lineage corresponds to the achieved minimum of the objective function.

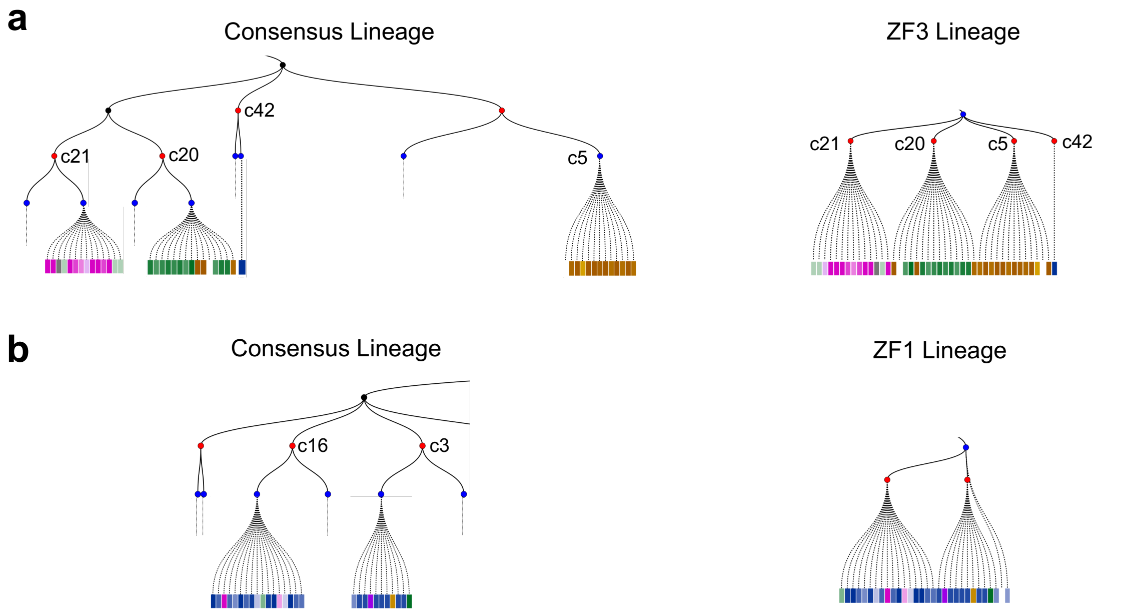

Supplementary Figure S5: Consensus lineage preserves ancestor-descendant relationships in individual lineages. (a) Clusters c20, c21, c5, c42 are present in the same subtree in both the consensus lineage and ZF3 lineage. (b) Clusters c16 and c3 are present in the same subtree in both the consensus lineage and ZF1 lineage.

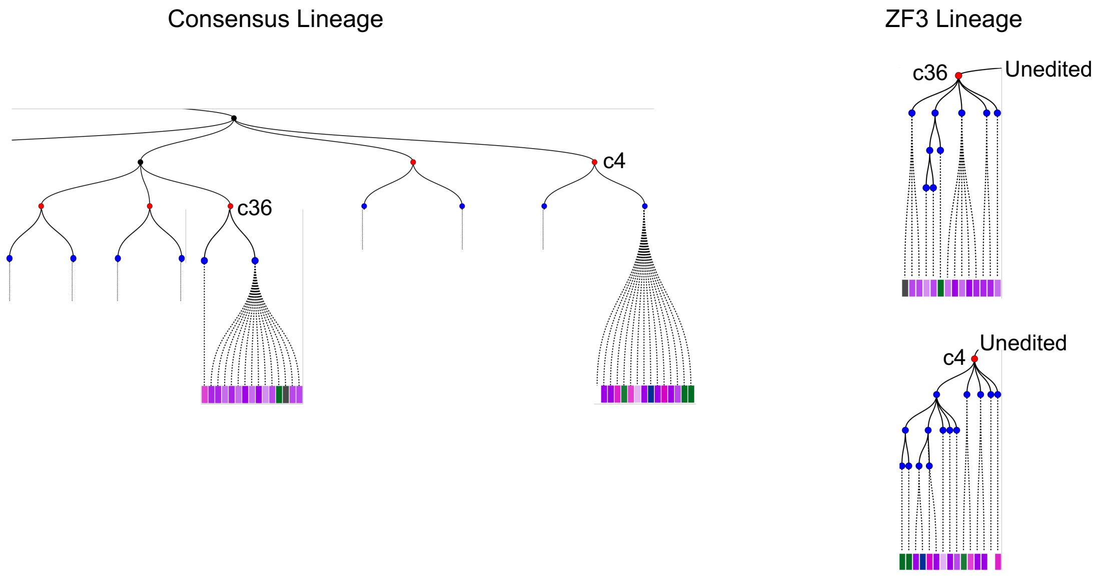

Supplementary Figure S6: Consensus lineage places similar cell clusters in the same subtree. In ZF3 lineage, clusters c36 and c4 both contain cells belonging to blood cell types but these clusters are placed in different branches from the root. In consensus lineage these clusters are placed in the same subtree.

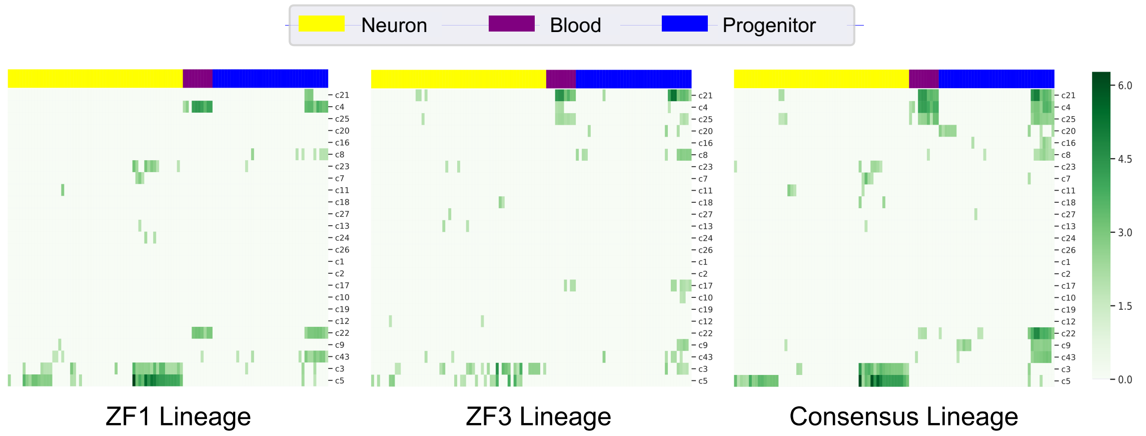

Supplementary Figure S7: Heat map of the p-values ($\sqrt{-\log(pvalue)}$, higher value means more significant) for all GO terms for the consensus clusters that contains 10 or more cells. Rows represent selected consensus clusters and the columns represent different GO terms (Supplementary Table S4). The values were colored as shown in the key. Yellow, purple and blue columns correspond to GO terms related to neurons, blood and progenitors respectively. The leftmost panel shows the heat map for ZF1, middle panel for ZF3 and the rightmost panel for the consensus lineage tree.

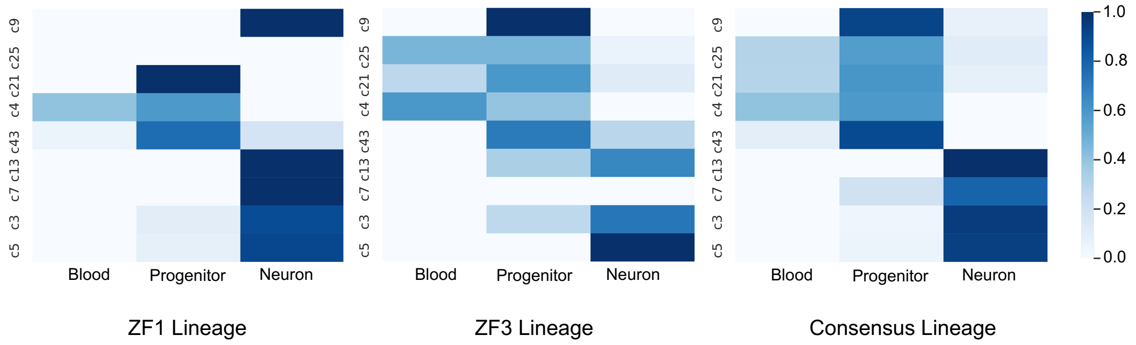

Supplementary Figure S8: Heat map of proportions of each type (neuron, blood and progenitor) of GO terms for selected consensus clusters. The rows represent selected consensus clusters and the columns represent different types of GO terms. Each row sums to 1.

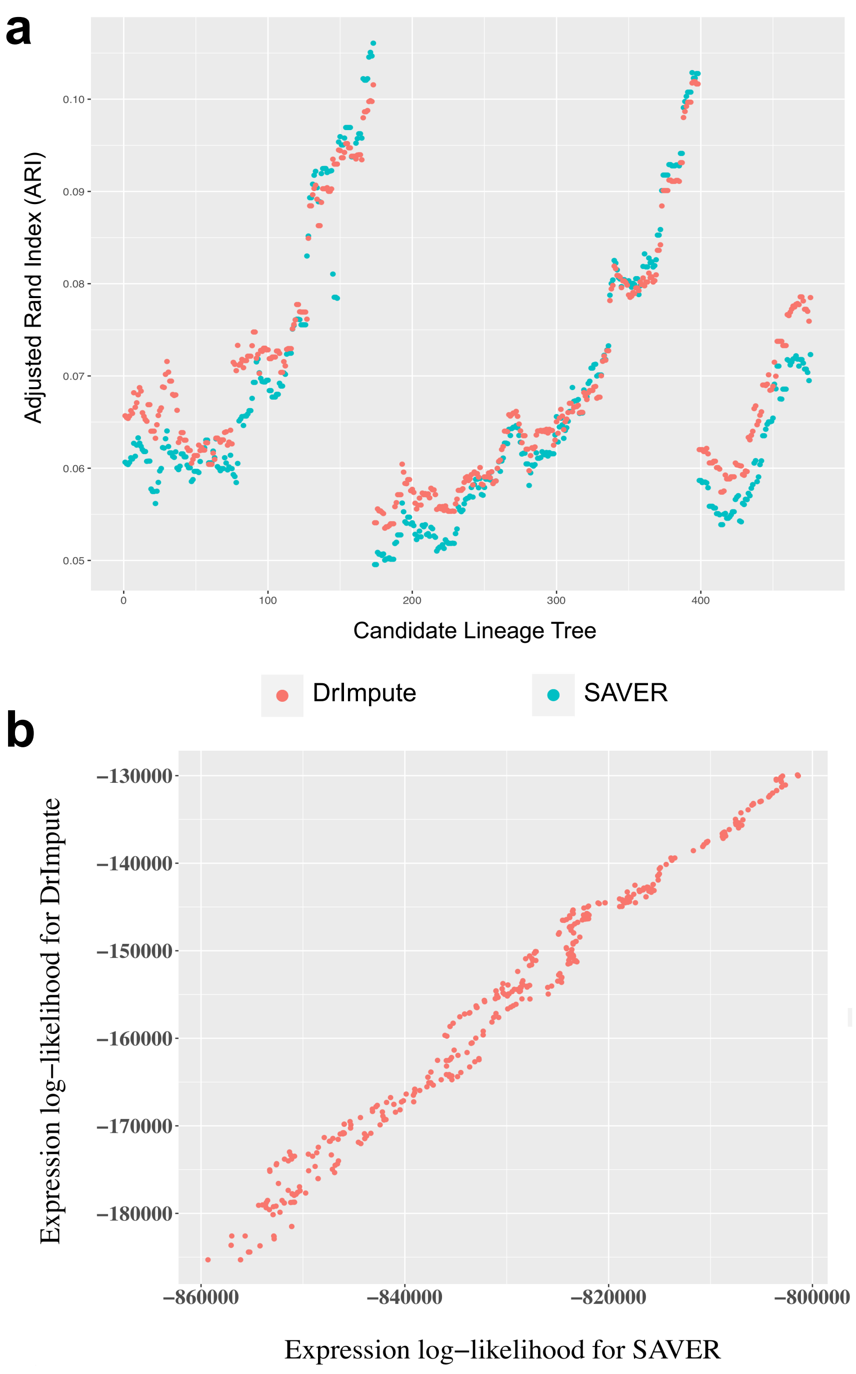

Supplementary Figure S9: (a) Effect of the imputation method on LinTIMaT's expression likelihood function displayed through cell clustering performance. For a set of candidate lineage trees for ZF3, we compared the cell clustering based on expression likelihood for expression data imputed using two imputation methods: DrImpute and SAVER. The cell clustering performance is measured in terms of Adjusted Rand Index. (b) Plot comparing the expression log-likelihoods for a set of lineage trees for data imputed using DrImpute and SAVER (correlation 0.9914).

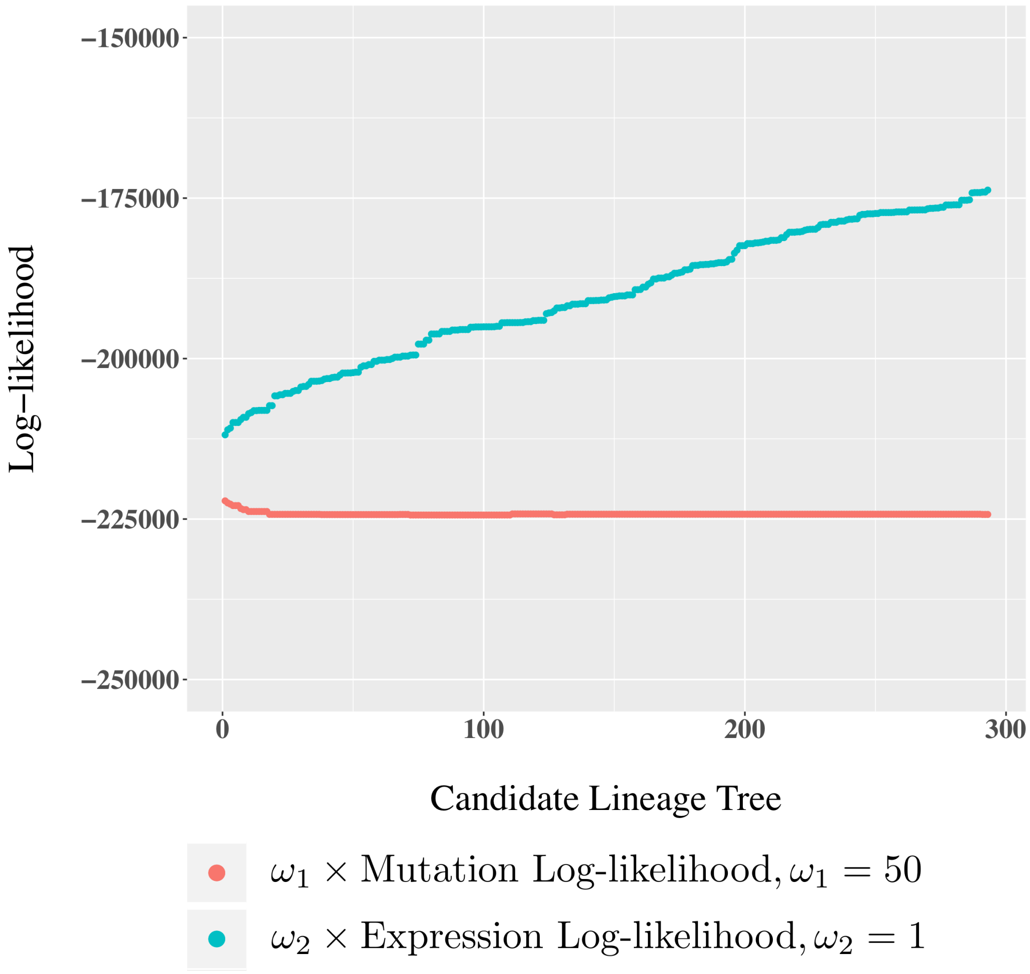

Supplementary Figure S10: Comparison of weighted values of mutation log-likelihood and expression log-likelihood for specific weights, $\omega_1 = 50$ and $\omega_2 = 1$ for a set of candidate lineage trees for ZF1. For these values of the weights, the weighted values of the two log-likelihood functions remain in the same range.

### 4 Supplementary Tables

Supplementary Table S1: The ARI for each scGESTALT tree calculated based on different levels

| ZF1 | | | ZF3 | | |
|---|---|---|---|---|---|
| Level | #cluster | ARI | Level | #cluster | ARI |
| 1 | 25 | 0.035658001 | 1 | 23 | 0.01938034 |
| 2 | 75 | 0.047930182 | 2 | 67 | 0.037909748 |
| 3 | 363 | 0.041749244 | 3 | 153 | 0.045092761 |
| 4 | 531 | 0.06103636 | 4 | 279 | 0.027522376 |
| 5 | 676 | 0.04716886 | 5 | 368 | 0.002327659 |
| 6 | 750 | 0 | 6 | 376 | 0 |
| Barcode | 192 | 0.061237582 | Barcode | 150 | 0.056133354 |

Supplementary Table S2: Comparison of log-likelihood score of lineage trees based on only mutation data

| Method | ZF1 | ZF3 |
|---|---|---|
| LinTIMaT | −4485.564873 | −2871.118661 |
| MP | −303463.075241 | −102474.127300 |

Supplementary Table S3: Strings for filtering the GO terms for each GO type

| **GO type** | **filter strings** |
|---|---|
| neuron | neuro nervous synap |
| blood | heme hema hemo erythrocyte myeloid hscs immune |
| progenitor | develop differentiat |

Supplementary Table S4: Full list of filtered GO terms used in GO p-value/proportion heat maps, see Supplementary Table S3 for the keywords we used to filter these GO terms.

| index | GO type | GO term | index | GO type | GO term |
|---|---|---|---|---|---|
| 1 | neuron | postsynaptic density | 47 | neuron | peripheral nervous system neuron axonogenesis |
| 2 | neuron | neuron to neuron synapse | 48 | neuron | regulation of trans-synaptic signaling |
| 3 | neuron | calcium ion-regulated exocytosis of neurotransmitter | 49 | neuron | signal release from synapse |
| 4 | neuron | chemical synaptic transmission | 50 | neuron | neuron differentiation |
| 5 | neuron | trans-synaptic signaling | 51 | neuron | Neuronal System |
| 6 | neuron | vesicle-mediated transport in synapse | 52 | neuron | neuron projection development |
| 7 | neuron | Presynaptic depolarization and calcium channel opening | 53 | neuron | anterograde trans-synaptic signaling |
| 8 | neuron | postsynapse | 54 | neuron | neuron projection cytoplasm |
| 9 | neuron | positive regulation of synaptic transmission, glutamatergic | 55 | blood | erythrocyte development |
| 10 | neuron | synaptic vesicle membrane | 56 | blood | erythrocyte homeostasis |
| 11 | neuron | Neurotransmitter release cycle | 57 | blood | hemopoiesis |
| 12 | neuron | positive regulation of synaptic transmission | 58 | blood | RUNX1 regulates transcription of genes involved in differentiation of HSCs |
| 13 | neuron | neurotransmitter secretion | 59 | blood | immune system development |
| 14 | neuron | neuron projection | 60 | blood | hematopoietic or lymphoid organ development |
| 15 | neuron | Glutamate Neurotransmitter Release Cycle | 61 | blood | myeloid cell differentiation |
| 16 | neuron | Acetylcholine Neurotransmitter Release Cycle | 62 | blood | myeloid cell homeostasis |
| 17 | neuron | postsynaptic specialization | 63 | blood | erythrocyte differentiation |
| 18 | neuron | Activation of NMDA receptors and postsynaptic events | 64 | blood | immune system process |
| 19 | neuron | presynaptic active zone | 65 | progenitor | multicellular organism development |
| 20 | neuron | synaptic vesicle transport | 66 | progenitor | epithelium development |
| 21 | neuron | neurogenesis | 67 | progenitor | nervous system development |
| 22 | neuron | synapse part | 68 | progenitor | cellular developmental process |
| 23 | neuron | synaptic signaling | 69 | progenitor | erythrocyte development |
| 24 | neuron | generation of neurons | 70 | progenitor | regulation of cell development |
| 25 | neuron | synapse | 71 | progenitor | embryo development ending in birth or egg hatching |
| 26 | neuron | establishment of synaptic vesicle localization | 72 | progenitor | anatomical structure development |
| 27 | neuron | regulation of neurotransmitter levels | 73 | progenitor | RUNX1 regulates transcription of genes involved in differentiation of HSCs |
| 28 | neuron | neurotransmitter receptor complex | 74 | progenitor | animal organ development |
| 29 | neuron | postsynaptic density membrane | 75 | progenitor | embryo development |
| 30 | neuron | neuron part | 76 | progenitor | cell development |
| 31 | neuron | synaptic vesicle cycle | 77 | progenitor | immune system development |
| 32 | neuron | peripheral nervous system neuron differentiation | 78 | progenitor | tissue development |
| 33 | neuron | synaptic vesicle localization | 79 | progenitor | peripheral nervous system neuron differentiation |
| 34 | neuron | presynapse | 80 | progenitor | hematopoietic or lymphoid organ development |
| 35 | neuron | postsynaptic specialization membrane | 81 | progenitor | peripheral nervous system neuron development |
| 36 | neuron | peripheral nervous system neuron development | 82 | progenitor | regulation of developmental process |
| 37 | neuron | synaptic vesicle exocytosis | 83 | progenitor | myeloid cell differentiation |
| 38 | neuron | synaptic vesicle | 84 | progenitor | cell differentiation |
| 39 | neuron | neuron development | 85 | progenitor | neuron development |
| 40 | neuron | modulation of chemical synaptic transmission | 86 | progenitor | developmental process |
| 41 | neuron | Neurotransmitter uptake and metabolism In glial cells | 87 | progenitor | regulation of cell differentiation |
| 42 | neuron | synaptic membrane | 88 | progenitor | system development |
| 43 | neuron | neurotransmitter transport | 89 | progenitor | neuron differentiation |
| 44 | neuron | asymmetric synapse | 90 | progenitor | neuron projection development |
| 45 | neuron | regulation of neurogenesis | 91 | progenitor | erythrocyte differentiation |
| 46 | neuron | Transmission across Chemical Synapses | 92 | progenitor | chordate embryonic development |

[6] Bushra Raj, Daniel E Wagner, Aaron McKenna, Shristi Pandey, Allon M Klein, Jay Shendure, James A Gagnon, and Alexander F Schier. Simultaneous single-cell profiling of lineages and cell types in the vertebrate brain. *Nature biotechnology*, 2018.
